# Near-perfect precise on-target editing of human hematopoietic stem and progenitor cells

**DOI:** 10.1101/2023.05.26.542436

**Authors:** Fanny-Meï Cloarec-Ung, Jamie Beaulieu, Arunan Suthananthan, Bernhard Lehnertz, Guy Sauvageau, Hilary M Sheppard, David JHF Knapp

## Abstract

Precision gene editing in primary hematopoietic stem and progenitor cells (HSPCs) would facilitate both curative treatments for monogenic disorders as well as disease modelling. Precise efficiencies even with the CRISPR/Cas system, however, remain limited. Through an optimization of guide RNA delivery, donor design, and additives, we have now obtained mean precise editing efficiencies >90% on primary cord blood HSCPs with minimal toxicity and without observed off-target editing. The main protocol modifications needed to achieve such high efficiencies were the addition of the DNA-PK inhibitor AZD7648, and the inclusion of spacer-breaking silent mutations in the donor in addition to mutations disrupting the PAM sequence. Critically, editing was even across the progenitor hierarchy, did not substantially distort the hierarchy or affect lineage outputs in colony-forming cell assays or the frequency of high self-renewal potential long-term culture initiating cells. As modelling of many diseases requires heterozygosity, we also demonstrated that the overall editing and zygosity can be tuned by adding in defined mixtures of mutant and wild-type donor. With these optimizations, editing at near-perfect efficiency can now be accomplished directly in human HSPCs. This will open new avenues in both therapeutic strategies and disease modelling.

## Introduction

Precise genome editing holds substantial promise for both accurate disease modelling and for potential curative treatments for monogenic disorders. There has been a particular interest in therapeutic editing in the hematopoietic system due to the ability to transplant cells to give life-long grafts, and the relatively large number of monogenic disorders that could be treated by gene repair/replacement. Precision edits can be generated using a CRISPR/Cas system to introduce a break at the target locus together with the addition of a template DNA to engage the homology-directed repair (HDR) pathway and insert the edit of interest^1^. These templates contain the change of interest flanked by homology arms matching the sequence on either side of the break site. While this strategy can function, non-homologous end joining (NHEJ) which results in random insertions and deletions (indels) is the default pathway, as HDR can only operate in the S and G2 phase of the cell cycle^2,3^. As such, the efficiency of HDR mediated editing, particularly in relevant primary human stem and progenitor populations remains limited, with most strategies achieving efficiencies in the range of 10-20%^4,5^. Despite such limited efficiencies, multiple clinical trials using CRISPR/Cas-based editing strategies are currently ongoing, reinforcing the extreme interest in the area^6^.

One simple method to improve HDR efficiency involves increasing the concentration of donor template as intracellular donor concentration has been shown to correlate to HDR efficiency^7^. Indeed, in HSPCs high concentrations of donor AAV have been shown to improve the efficiency of HDR, though likely at the cost of cell viability as, while not measured directly, a subsequent live-dead selection was performed in the study^8^. As cell numbers are limiting and functionality is critical, editing toxicity is as important as raw efficiency. To this end, transient knockdown of p53 by siRNA was shown to improve survival of human hematopoietic stem cells (HSC), though not directly modulate the HDR editing efficiency^9,10^. Interestingly, Inhibition of 53BP1 which has the additional effect of NHEJ inhibition has been shown to improve the editing efficiency in these cells reaching up to ∼60%^10^. Critically, both studies demonstrate that edited cells retained their functional stem cell capacity (long-term engraftment in xenotransplantation) following the editing protocol, with efficiencies similar to pre-transplant measurements. Another small molecule inhibitor of NHEJ, M3814 that inhibits DNA-PK, has been shown to increase editing efficiencies up to 80-90% in primary human T cells^11^. M3814 also has the benefit of being a small molecule rather than protein like the inhibitor of 53BP1, thus making it cheaper and easier to use. Importantly for safety, the 53BP1 study also showed that while NHEJ inhibition increased the rate of large deletions in the absence of HDR donor, when HDR donor was present the rate of large deletions was not increased, nor was the rate of off-target edits^10^. Inhibition of NHEJ is thus likely a safe and effective way to improve the rates of HDR.

In this study, we optimized ribonuclear protein (RNP) concentration and selection, donor type and design, and small molecule additives, and combined this with optimal culture time and conditions needed to induce division while maintaining stemness in primary human HSC^12,13^ in order to further refine HDR in these cells. We demonstrate that while M3814 increases HDR editing efficiency in human HSPCs, a more specific DNA-PK inhibitor AZD7648^14^ is capable of further improving this efficiency. Combining AZD7648 with optimal pre-stimulation, RNP, p53 siRNA and AAV donor concentrations yielded mean efficiencies of 97% editing with minimal toxicities. Surprisingly, these conditions worked at equivalent efficiencies for short single-stranded oligodeoxynucleotide donors (ssODNs) giving mean efficiencies of 94% when the ssODNs were modified to mutate not just the PAM sequence but also multiple positions in the spacer (silent mutations). These donors which thus far have not previously been effective in HSPC are much faster and easier to design and are fully synthetic, thus facilitating rapid prototyping and downstream translation. Importantly, the editing protocol is consistent across the hematopoietic hierarchy, and does not affect lineage choice in CFC assays or high self-renewal potential long-term culture initiating cell frequency. To further facilitate disease modelling applications where zygosity is an important consideration, we also demonstrated that zygosity can be tuned by providing a mixture of silent and mutant donors. Our refined protocol will thus facilitate both therapeutic editing strategies and disease modelling strategies.

## Methods

### Human cord blood processing and cryopreservation

Anonymized consented human umbilical cord blood was obtained from Hôpital St-Justine and Hema Quebec, Montréal, QC, Canada. Ethics approval for the use of these cells was granted from the Comité d’éthique de la recherche clinique (CERC) of the Université de Montréal. CD34+ cells were isolated using EasySep™ Human CD34 Positive Selection Kit II (STEMCELL Technologies, Vancouver, BC, Canada) as per manufacturer instructions, and then used either directly or cryopreserved in fetal bovine serum (Gibco) and 10% dimethyl sulfoxide (BioShop). Cells were frozen slowly at -80 °C in a CoolCell (Corning) and transferred the following day to liquid nitrogen until use. On thaw, cells were rapidly warmed to 37 °C in a water bath, diluted 10x in RPMI + 10% FBS (Gibco) and spun to remove media and residual DMSO. Viable cells were counted using a hemocytometer with Trypan Blue (Gibco) and placed into culture.

### CD34+ cell culture

CD34+ cells were cultivated in StemSpan™ SFEM II media supplemented with 1X StemSpan™ CC100 and 35 nM UM171. Cell density was maintained below 250000 cells/mL of total media to prevent autoinhibition of the primitive cells^12,15^. Cells were maintained in a humidified incubator at 37 °C with periodic viable cell counts (by hemocytometer) to ensure that the density remained within the acceptable range.

### RNP assembly

The tracrRNA and crRNA (IDT) were mixed at equimolar ratio to a final concentration of 100 μM, annealed for 5min at 95 °C and cooled to 25 °C at 0.1 °C/s. Annealed gRNA was then added to Cas9 enzyme (IDT) and incubated at room temperature for 15 min with a ratio of Cas enzyme to gRNA of 1:2.5.

### CD34+ cell editing

After 48 hour of pre-stimulation, viable cells were counted by hemocytometer. Prior to nucleofection, cells were washed once with PBS, spun down for 5 minutes at 300 g, and re-suspended in buffer 1M^16^ (5mM KCl; 15mM MgCl2; 120mM Na2HPO4/NaH2PO4 pH7.2; 50mM Manitol) such that each well of the Nucleocuvette strip would contain 20 000 – 100 000 cells. Assembled RNP, p53 siRNA (20 fmol, Thermo id s605), electroporation enhancer (IDT), and any ssODN donors (IDT) (as specified in for each experiment). Overall RNP and other additives were kept at or below 10% of the total 20 μL volume per well. Handling time between wash and nucleofection was kept within a 10-minute window. Cells were nucleofected using the Lonza 4D nucleofector device with nucleocuvette strips, Primary P3 and DZ100 program. Following nucleofection, cells were allowed to rest for 5 minutes then added to pre-warmed wells of a 24 well plate containing media (as specified in CD34+ cell culture) supplemented with small molecules as indicated for specific experiments (AZD7648, RS-1, Cayman Chemicals; M-3814, Toronto Research Chemicals). Where AAV donor was used, it was added to the well within 15 minutes of nucleofection^5^. Custom AAV donors were generated by the Canadian Neurophotonics Platform Viral Vector Core Facility (RRID:SCR_016477). Cells were incubated for an additional 48 hours prior to subsequent use. After this 48-hour period, viable cells were counted by hemocytometer with Trypan Blue.

### Genomic DNA lysis and amplification

At time of harvest cells were centrifuged at 300g for 5 min and washed once with PBS, and pelleted again followed by re-suspension in a gDNA lysis buffer (50 mM Tris, 1 mM EDTA, 0.5% Tween-20 and 16 U/mL Proteinase K). Samples were incubated 1 hour at 37°C and 10 minutes at 95°C. Lysates were amplified by PCR using the Platinum Taq SuperFi II Master Mix with 0.5 μM of each primer, and lysate comprising no more than 5% of the total volume of the reaction. For T7E1 assays, short, and long ssODN without silent mutations, a single amplification was performed with primers pri0002-H3 + pri0002-H4 (Supplementary Methods). For short ssODN with silent mutations a single amplification was performed with pri0077-F + pri0077-R (Supplementary methods). For AAV, a nested PCR was performed with where the first PCR was performed (SRSF2= pri0077-F + pri0077-R, SF3B1= pri0078-F + pri0078-R; Supplementary methods) followed by purification using a GeneJet PCR purification kit (Thermo) and 15-30 ng transferred to a second PCR (SRSF2= pri0002-H3 + pri0002-H4, SF3B1= pri0002-H1 + pri0002-H2; Supplementary methods). For analysis of off-target cutting, the top 3 sites predicted in Benchling were amplified with primers as indicated in the Supplementary Methods, and sequenced using either indicated sequencing primers (off-targets 2 & 3) or one of the amplification primers (off-target 1). For all PCRs, the program was 98°C 30 s; 35× (98°C 10 s, 60°C, 10 s, 72°C 30 s); 72°C 5 min.

### Measurement of RNP cutting efficiency after 48h post-electroporation using the T7E1 assay

PCR products were purified using the GeneJet PCR purification kit, and DNA quantified by nanodrop. 100 ng of DNA was added to 1x NEBuffer 2, and annealed as follows: 95°C 5 min; 95-85°C -2°C/s; 85-25°C -0.1°C/sec; hold at 4°C. The sample was then split in half, with half kept aside as a non-digested control, and 2.5 U T7 Endonuclease I added to the other half. All samples are incubated at 37°C for 15 minutes and then run on a 2% TAE agarose gel with GelRed (Biotium). Gels were imaged at non-saturating intensities using a GeneGenius Imaging System (Syngene). The ratio of cut to uncut bands was then quantified by densitometry.

### Quantification of HDR donor integration

HDR donor integration was assessed by three different ways depending on the HDR donor. For the PAM mutant only short and long ssODNs, a BspEI site was introduced by the mutation. For these, 100 ng of amplified DNA was added to 1x NEBuffer 3.1 with 2 U BspEI. These were then incubated at 37 °C for 1 hour then run on a 2% TAE agarose gel with GelRed and quantified by densitometry. For AAV donors, these were run directly on a gel and integration quantified based on the size shift from the introduced 143 bp synthetic intron. Finally, for ssODN with silent mutations, PCR products were purified by GeneJet PCR purification, quantified by nanodrop, and 5-15 ng sent for Sanger sequencing at the IRIC Genomics core with the primer pri0003-A1 (Supplementary methods). Integration was then quantified using ICE Analysis (Synthego Performance Analysis, 2019. v3.0. Synthego) with comparison between each edited sample back to a matched unedited control.

### Analysis of off-target editing by nanopore sequencing

Amplicons for each of the three predicted off-target sites were first pooled per sample at equimolar ratios based on their known length and concentration measured by Nanodrop. This allows higher multiplexing without requiring additional barcoding reagents as each amplicon will map uniquely. Each off-target pool was then end repaired using the NEBNext® Ultra™ II End Repair/dA-Tailing Module with the addition of DNA Control Sample (Oxford Nanopore) as per manufacturer instructions, and end-repaired/A-tailed products purified using AmpureXP beads at a 1x bead ratio. Unique native barcodes were then added to each repaired/tailed amplicon pool using the Native Barcoding Kit 24 V14 (Oxford Nanopore) and NEB Blunt/TA Ligase Master Mix as per manufacturer instructions, all barcoded amplicons pooled into a single tube, and the library purified using AmpureXP beads, this time at a 0.7x bead ratio. Finally, sequencing adapters were added to the pooled library using the NEBNext® Quick Ligation Module (NEB) with the native adapter from the Native Barcoding Kit 24 V14, and this again purified with AmpureXP beads using Short Fragment Buffer (Oxford Nanopore) instead of 80% ethanol, all as per manufacturer instructions. Final concentration was determined by Nanodrop and 20 fmol loaded onto a Flongle Flow Cell (R10.4.1) with a minION sequencing device (MIN-101B) with Flongle adapter (all from Oxford Nanopore). Samples were run with MinKNOW, and base-calling executed on Super-Accurate mode.

Following base-calling, a custom pipeline based on bcftools was used to call variants (available at: https://github.com/djhfknapp/Nanopore_Amplicon_CRISPR_Analysis). For this, fastq files were first pooled per barcode using Biopython. These were then aligned to the known amplicon sequences (from GRCh38) using minimap2 with the map-ont option, converted to a BAM file, sorted, and indexed using samtools view, sort, and index respectively. Variants were then called using bcftools mpileup with options ‘-Ov -Q 16 -a FORMAT/DP,FORMAT/AD -d 10000 --min-MQ 10’. Next a per site summary was generated from the VCF file counting all high-quality (those meeting the bcftools criteria, ie. base quality ≥16, mapping quality ≥10) reference reads, substitutions, deletions, and insertions, and output as a TSV file. This output was used directly for per-sample plotting. For summary statistics, the predicted target sites were identified for each amplicon, and the percentage of reads supporting specific substitutions, deletions (starting in or extending into the target site), or insertions were calculated per site. Only variants supported by at least 3 reads and making up at least 0.1% of total reads at a site were counted for these summary statistics. Assuming that any given read has only a single variant which impacts the target region, fully reference percentages were then calculated as 100 minus the sum of all substitutions, deletions, and insertions. This assumption was generally valid with the target sequences used and if anything would under-represent percent of fully reference matching reads.

### Gel Densitometry

Band quantification was performed using GelAnalyzer 19.1 (www.gelanalyzer.com). Briefly, lanes were detected using the detect bands on every lane function. Peaks were then detected automatically using the “Detect Bands on Every Lane” function. Background was subtracted using the Rolling Ball option within autodetect background on all lanes with a peak width tolerance of 15%. The contribution of each band was then calculated as the density of that peak divided by the total density of all peaks.

### Flow cytometry and sorting

48 hours post-edit, cells were collected, spun down, and stained in PBS + 2% FBS using a panel of 4 antibodies: CD34 (1:200, AF647 Mouse Anti-Human CD34 (clone 581)), CD45RA (1:100, V450 Mouse Anti-Human CD45RA (clone HI100)), CD90 (1:200, PE-CF594 Mouse Anti-Human CD90 (clone 5E10)) and CD49c (1:50, FITC anti-human, CD49c (Clone REA360)) for 1h on ice in the dark. Precision Count Beads (BioLegend) were added prior to sort to allow quantification of absolute numbers. Cells were analyzed and sorted on a BD FACSAria III sorter. Cells were either sorted for CD34+ for colony-forming cell (CFC) assays, or for 4 sub-populations as follows: long-term HSCs (LT-HSC; CD34+CD45RA-CD90+CD49c+), Intermediate HSCs (IT-HSC; CD34+CD45RA-CD90+CD49c-), multi-potent and erythroid progenitors (MPP/E; CD34+CD45RA-CD90-CD49c-) and late progenitors (Adv-P; CD34+CD45RA+). In all cases this was done on purity mode using dilute samples to minimize cell lose and prevent possible contamination. FACS data was analysed using FlowJo software (version 10.8.1).

### CFC assays

For CFC assays, 500 and 1500 CD34+ cells were added to 1 mL of MethoCult™ H4034 Optimum medium (StemCell Technologies) and placed into one well of a 6-well plate. Plates were incubated for 14-16 days in a humidified incubator at 37°C, 5% CO_2_. Wells were then imaged in colour using a Cytation 5 (BioTek), and colonies scored based on colour, morphology and size as previously performed^17^. Selected colonies were then picked into PBS, spun down, and subjected to gDNA extraction, PCR and integration assessment as detailed above.

### Long-Term Culture Initiating Cell (LTC-IC) Assays

Cells were either edited with silent SRSF2 ssODN donors as per our optimal protocol: 48-hour pre-stimulation in StemSpan + CC100 + 35 nM UM171, delivery of editing components by nucleofection of 30.5 pmol RNP + 50 pmol ssODN + 20 fmol p53siRNA, and 48-hour culture in StemSpan + CC100 + 35 nM UM171 + 0.5 μM AZD7648, or controls maintained in StemSpan + CC100 + 35 nM UM171 for the 4 day period. At the end of this period, single LT-HSC (CD34+CD45RA-CD90+CD49c+) were sorted per well into the inner 60 wells of a flat-bottom 96 well LTC-IC plate. Edited cells were sorted into half of each plate and unedited controls into the other half. LTC-IC assays were performed as described in ^18^. Briefly, one day before sorting, each well of the 96 well plate was first coated with 45 μL 2.25% Type I Bovine Collagen Solution (StemCell Technologies) for 1 hour at 37 C. Next, 5×10^4^ cells comprising an equal mixture of irradiated M210B4 fibroblasts expressing human IL-3 and G-CSF, sl/sl mouse fibroblasts expressing human SCF and IL3, and sl/sl mouse fibroblasts expressing human FLT3L were added to each well. The day of the sort, media was changed to 100 μL Myelocult H5100 (StemCell Technologies) supplemented with 10^−6^ M hydrocortisone (BioShop). Following the sort, cultures were maintained at 37°C in a humidified incubator with 5% CO2 with weekly half-media changes for 6 weeks. Each week wells were imaged on a Cytation 5 imager, and scored for the presence of refractile non-adherent cells. At the end of 6 weeks, cultures were supplemented with 50 ng/mL recombinant human SCF + 20 ng/mL each of GM-CSF, IL-3, IL-6, G-CSF, and 3 U/mL erythropoietin (all from Genscript). Cultures were allowed to continue for an additional 2 weeks and then imaged and scored again. For this final timepoint, confluent cultures were scored as highly proliferative, those with >50 cells but below confluence were scored as low proliferation, those which had detectable cells at earlier time points but no longer at the final scoring were scored as transient, and those in which no clones were ever detected were scored as negative.

### Single cell cloning for zygosity analyses

Single CD34+CD45RA-cells were sorted (see staining and sort details above) 48h post-editing on single cell mode into independent wells of a 96-well plate and cultured as detailed in CD34+ cell culture for up to 14 days. When a clone reached a large size (100s-1000s of cells) they were harvested, or at the end of the 14-day period those with visible but small clones (<100 cells) were harvested. In each case these underwent lysis, PCR and gel assessment. All integration negative colonies were assessed for silent donor integration by Sanger sequencing/ICE analysis as described above.

### Statistical analyses

All statistical analyses were performed in R (version 4.1.2). For statistical comparisons, paired t-tests were performed in cases where the same cord was used for all conditions and unpaired where necessary. In all cases, an FDR correction for multiple testing was performed using the ‘p.adjust’ function. For LTC-IC frequencies, Extreme Limiting Dilution Analysis was performed using R function ‘elda’ from the package ‘statmod’.

## Results

### Precise editing can be achieved using both AAV and ssODN donors in cord blood CD34+ cells

As a basis for our protocol (Figure 1A), we used a 48-hour pre-stimulation in a growth factor mixture of SCF, FLT3L, IL3, and IL6 which we have previously shown induces even the most primitive phenotypic HSC to cycle by ∼66 hours^12^, thus ensuring that editing machinery would be present during S/G2 phase of the cells. We also included UM171, which improves the maintenance of HSC functional capacity^13^ and can improve transduction efficiency in these cells^19^. These conditions are similar to other reported editing strategies^4^. As a first step in our optimization, we tested different conditions for gRNA-mediated cutting. We tested two concentrations of RNP with and without an electroporation enhancer (IDT) on a site in the SRSF2 locus (Figure S1). Cutting efficiency was assessed 48h post electroporation using the T7E1 assay^20^ and toxicity by counting the absolute cell number for each condition on the same day (Figure S1). At lower RNP concentrations, we observed a slight benefit of the addition of electroporation enhancer, though this was not present at higher concentrations (FDR=0.02 and FDR=0.48 respectively, Figure S1A). Similarly, increasing RNP concentration also showed a benefit to overall editing efficiency (FDR=0.08, Figure S1A). Both the addition of electroporation enhancer and the increase in RNP concentration, however, resulted in significantly decreased cell numbers indicating toxicity (p=0.003, Figure S1B). Similar patterns were observed at the SF3B1 locus, though overall efficiency was lower for this gRNA (Figure S1C,D). As the improvements in cutting efficiency, while statistically significant, were only in the range of ∼10% and both increased toxicity, we selected 30.5 pmol RNP without enhancer for subsequent experiments.

**Figure 1.**
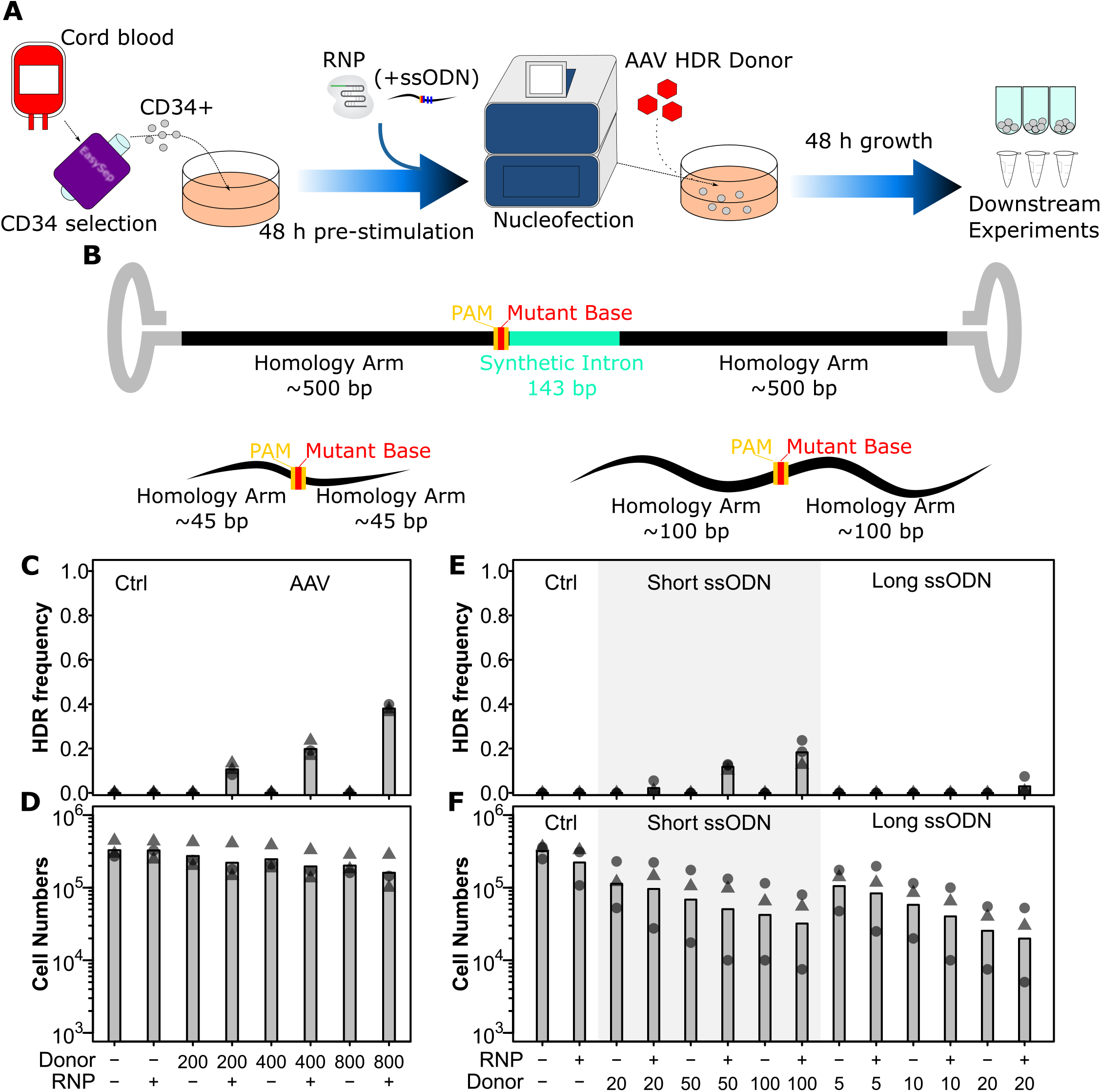
AAV and short ssODN both allow precise editing in human HSPCs. **A)** *Experimental design*. **B)** *HDR donor configurations*. The SRSF2 P95H AAV donor is shown above and short and long ssODN donors are shown below with features indicated. Annotated sequences are shown in Supplementary Methods. **C)** *HDR integration efficiency by AAV dose*. Cells were edited with 30.5 pmol RNP (or not as indicated) with indicated multiplicities of infection (MOI) of AAV donor. Bars show mean values and points show measurements for individual cords. Male cords are shown as triangles and female as circles. **D)** *Viable cell number by AAV dose*. Hemocytometer counts at the time of harvest are shown for each sample from (**C**). **E)** *HDR integration efficiency for short and long ssODN donors*. Donor DNA amounts are shown in pmol. **F)** *Viable cell number by ssODN dose*. False-discovery rate (FDR) corrected paired t-test significance values are shown in Table S1.

We next tested three different types of donors: AAV, short ssODN, and long ssODN (Figure 1B). Both ssODN donors were co-delivered in the same nucleofection as the RNP, while the AAV donors were delivered within 15 minutes of the electroporation as reported by Charlesworth et al^5^. Integration was assessed 48 hours later by PCR and gel quantification. For the AAV donor, this could be done by assessing product size directly as it included an ∼100bp synthetic intron. One critical factor we discovered here was that for this assessment a nested PCR was required with outer primers outside the homology arms of the AAV, as otherwise non-specific amplification from non-integrated donors predominated (Figure S2A). For ssODN donors, a BspEI site was introduced, allowing assessment based on an enzymatic digest. Using defined mixtures of WT and BspEI DNA, we show that this assay is accurate down to ∼1% integration efficiency (Figure S2B). As with RNP, we also assessed cell toxicity by absolute counts at the time of harvest. For AAV, a multiplicity of infection (MOI) of 400 MOI gave the highest integration rate prior to observable toxicity with an integration rate of 19.8% (Figure 1C,D). This integration is consistent with what can be found in the literature for this type of donor^4,5,9^. While the long ssODN donors showed a very low integration efficiency of 3% at 20 pmol and a high toxicity, short ssODN donors were more promising with an integration rate of 12% at 50 pmol and a lesser though still appreciable toxicity (Figure 1E,F). Higher concentrations of short ssODN further raised efficiency though with a corresponding increase in toxicity beyond a usable range (Figure 1E,F). Interestingly, toxicity for both ssODN donors was independent of the presence of the Cas9 nuclease (equivalent in the no RNP condition), suggesting that it is likely innate immune reaction to the single stranded DNA rather than the editing process itself. These data demonstrate that the AAV donor at 400 MOI and short ssODN at 50 pmol are the most suitable donors.

### Optimal inhibition of NHEJ enables integration efficiencies up to 100% for with both AAV and ssODN donors

To further increase HDR efficiency, we next tested whether the DNA-PK inhibitor M3814 could also boost HDR efficiency in HSPCs. In these tests, AAV donors were used and p53 siRNA included to maximize survival. We observed an increased HDR efficiency up to 70% with M3814 (Figure 2A), equivalent to reports with inhibition of 53BP1^10^. We also tested another DNA-PK inhibitor AZD7648 which has been reported to inhibit DNA-PK with reduced off-target activity on PI3K^14^. Interestingly, we observed improved editing efficiency at 0.5 μM with AZD7648 compared to M3814, and 0.5 μM AZD7648 was equivalent to 5 μM M3814 (Figure 2A, FDR=0.01 and FDR=0.24 respectively). Toxicity, as assessed by viable cell counts, was also marginally but statistically significantly lower for AZD7648 compared of M3814 at 0.5 μM (Figure 2B, FDR=0.02 for 0.5 μM). There was also no significant difference in editing efficiency for AZD7648 at 0.5 and 5 μM but a slight improvement in viability (FDR=0.16 and FDR=0.01 respectively Figure 2A,B). We next tested whether the addition of RS1, a RAD51 stabilizer and thus HDR enhancer could further improve efficiencies^21^. In this context the addition of RS1 either alone or together with AZD7648 was detrimental to editing efficiency and toxic to cells (Figure 2C). The combination of p53siRNA and AZD7648, however, was able to achieve mean efficiencies of 80% and up to 96% in these tests with minimal toxicity (Figure 2C,D). Repeating these tests on the SF3B1 locus revealed similar trends, though with a somewhat lower mean editing efficiency of 57% (Figure S3A,B). This difference reflects the lower cutting efficiency observed at this locus (Figure S1C,D). A closer examination of experimental factors that may affect efficiency revealed this to be related to the number of conditions tested in each experiment (Figure 2E). This suggests that there are likely technical factors associated with cell handling and nucleofection timing that play an important role in overall efficiency. Overall, these data suggest that limiting the number of co-processed samples, and the addition of 0.5 μM AZD7648 for the 48 hours following editing could substantially improve editing to near perfect efficiencies.

**Figure 2.**
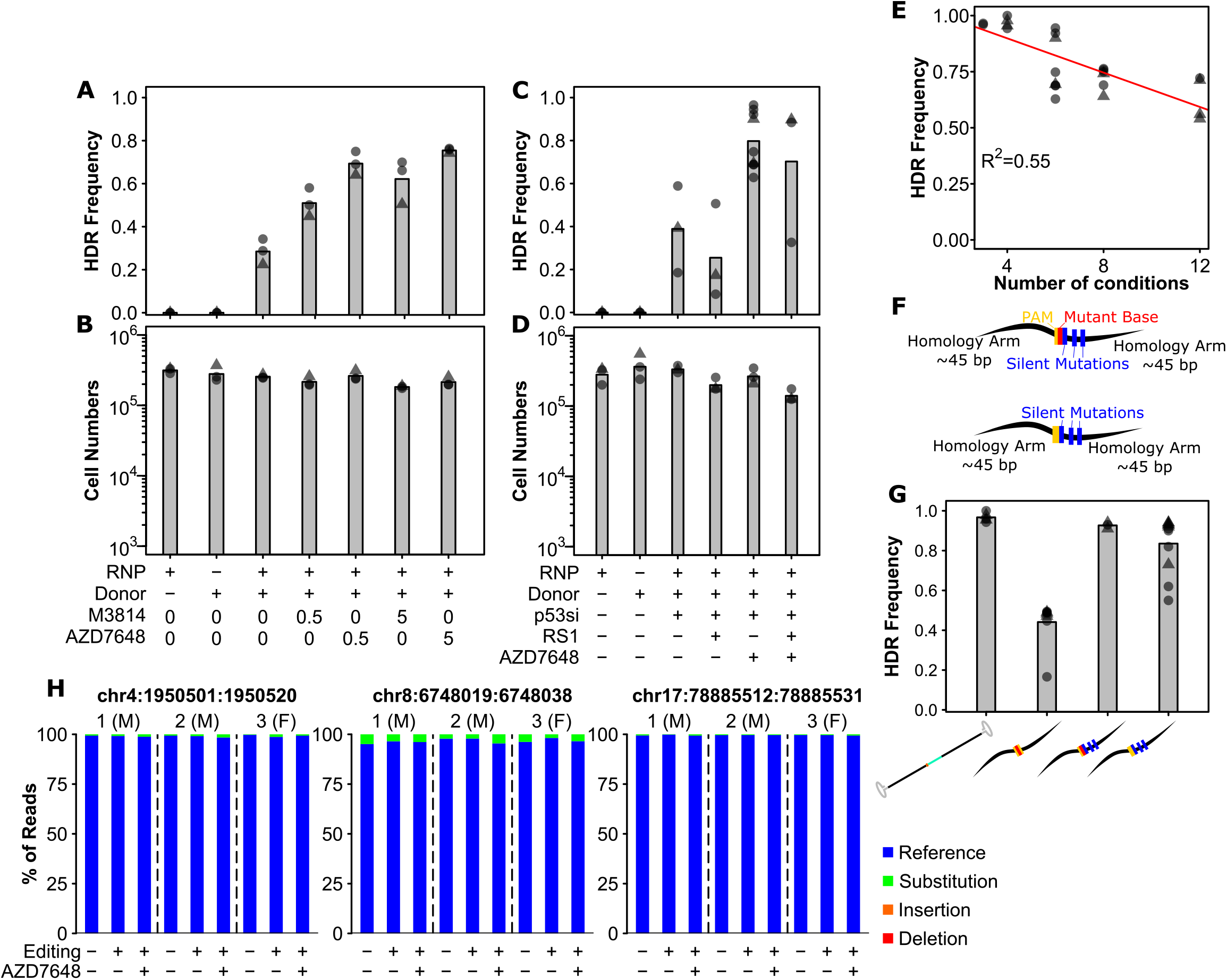
Small molecule-mediated inhibition of DNA-PK and optimal donor design substantially improve precise editing efficiency. **A)** *AZD7648 and M3814 improve HDR efficiency in primary human HSPC*. Cells were edited with 30.5 pmol RNP (or not as indicated) with 400 MOI of AAV donor and small molecules added as indicated (in μM). Bars show mean values and points show measurements for individual cords. Male cords are shown as triangles and female as circles. B) *Viable cell numbers with AZD7648 and M3814 addition*. Hemocytometer counts at the time of harvest are shown for each sample from (**A**). **C)** *HDR efficiency with combinations of AZD7648, p53 siRNA and RS-1*. Cells were edited with 30.5 pmol RNP (or not as indicated) with 400 MOI of AAV donor in the presence of the indicated additives. AZD7648 was used at 5 μM, p53 siRNA at 20 fmol, and RS-1 at 15 μM. D) *Viable cell numbers with additive combinations*. Hemocytometer counts at the time of harvest are shown for each sample from (**C**). **E)** *Technical factors associated with high sample number is associated with decreased HDR efficiency*. HDR efficiency is shown for all 30.5 pmol RNP, 400 MOI AAV, 20 fmol p53 siRNA, and 5 μM AZD7648 samples by the number of conditions processed in a given experiment. A linear fit is indicated as a red line. The R^2^ is indicated, and overall p value was <<0.001. **F)** Alternative designs for ssODN donors with key features indicated. Annotated sequences are shown in Supplementary Information. **G)** *Silent mutations allow ssODN donors to achieve similar efficiencies to AAV*. All edits were performed with 0.5 μM AZD7648, 20 fmol p53 siRNA, 50 pmol ssODN or 400 MOI AAV as indicated. Donor types are shown as their logos from (**1B, 2G**). **H)** *No observable off-target mutations at predicted target sites even with the addition of AZD7648*. The overall percent of reads containing exclusively reference allele, or any substitutions, deletions, or insertions that overlap with the predicted off-target cut sites is shown for 3 individual cords across the top 3 cut sites. Cells from each individual cord were split into an unedited control, and cells edited with the silent mutation containing ssODN for the SRSF2 locus under either standard conditions (ie. no p53siRNA or AZD7648) or with our optimal editing protocol (ie. with p53siRNA and 0.5 μM AZD7648). False-discovery rate (FDR) corrected paired t-test significance values are shown in Table S1.

With our handling and additive conditions, we next tested whether ssODN could be brought to similar efficiencies. Confirming our observed negative correlation between condition number and efficiency, in these tests performed with no more than 4 simultaneous conditions at nucleofection, we observed a mean efficiency of 97% with AAV donors (Figure 2F,G). With the short ssODN we observed a mean editing efficiency of 44% under these conditions, increased from the 13% but still substantially lower than AAV (Figure 2F,G), and consistent with reports for 53BP1 inhibition^10^. Further thought into the differences between the two suggested that perhaps it was the further disruption of the spacer sequence provided by the synthetic intron in the AAV donor which provided the difference (Figure 2F). As such we redesigned the ssODN donor to incorporate silent mutations throughout the PAM proximal bases of the spacer (Figure 2F). We also designed an equivalent donor which did not mutate the PAM but only contained the spacer mutations (Figure 2F). Interestingly, the incorporation of these additional spacer-breaking mutations drastically improved editing efficiency up to 94%, nearly that of the AAV (Figure 2G, S2C). Surprisingly, even the design incorporating only the silent spacer mutations, but no PAM mutation achieved a mean efficiency of 84%, further highlighting the importance of spacer mutations in donor design. Importantly, this latter design also contained only silent mutations, selected for codons with equivalent frequency to wild-type, and thus should have no effect on the biology of cells carrying them. Finally, as our protocol temporarily inhibits NHEJ and DNA damage surveillance, we tested whether cells edited under optimal conditions showed an increased off-target editing or not. Analysis of the top 3 predicted off-target sites in 3 sets of samples revealed no detectable edits at these sites either with or without AZD7648 (Figure 2H, S4, S5). Some substitutions were present both in and around the predicted off-target regions, however these were equivalent for a given individual between both edited conditions (+/-AZD7568) and unmanipulated controls (paired t-test FDRs >0.3, individual values in Table S1) indicating they were not introduced by the editing process. This suggests that off-target editing at predicted sites is not affected by AZD7648 addition, at least with well-designed gRNA. These modifications to donor design allowed even ssODN to reach near-perfect efficiency in primary human HSPCs.

### Editing efficiency is even across the hematopoietic hierarchy and does not disrupt phenotype proportions or self-renewal and differentiation functions

We next wanted to determine whether editing efficiency was equivalent across the hematopoietic hierarchy, as long-term HSC (LT-HSC) are a minority cell type within the overall CD34+ compartment. While phenotypes involving CD49f are highly selective on fresh cells^12,17,22^, not all of these markers are stable on cultured cells. As such, we used a panel consisting of only culture-stable markers which retained the ability to sort sub-populations from across the hierarchy. Populations sorted included LT-HSCs (CD34+CD45RA-CD90+CD49c+), intermediate HSCs (IT-HSC; CD34+CD45RA-CD90+CD49c-), multipotent and erythroid progenitors (MPP/E; CD34+CD45RA-CD90-CD49c-) and mature progenitors (Adv-P; CD34+CD45RA+)^23^. Of note, the PAM mutation in our mutant donors introduces a P95H mutation which is associated with myelodysplastic syndrome (MDS), acute myeloid leukemia (AML) and clonal hematopoiesis^24–26^. While this mutation is of interest in future disease modelling, to isolate the effects of the editing process itself on HSPC phenotype and function, biological assays were performed exclusively with silent donor. We observed no significant differences in editing efficiencies from bulk measurements regardless of sub-population suggesting that the editing was even across the progenitor hierarchy (Figure 3A, S6). We also observed a generally equivalent representation of each population within the CD34 compartment between silent edited and control cells, though a slight but statistically significant decrease was observed in the proportion of CD34+ cells of the advanced progenitor phenotype (FDR= 0.007; Figure 3B). To test whether these cells retain proliferative and neutrophil/monocyte/erythroid differentiation potential, we next performed colony-forming cell assays on edited and control cells (Figure 3C). Consistent with our earlier observations that donor alone induced some toxicity, we observed a slight drop in colony number even without editing (Figure 3D). Colony number was further decreased with editing (Figure 3D). This is consistent with the slight drop in the advanced progenitor phenotype which contains primarily neutrophil/monocyte producing progenitors. Importantly, however, no changes to colony type were observed (Figure 3E), suggesting that while there is a quantitative drop in progenitor function, this was even across progenitor types and the capability of remaining cells was not affected. Analysis of the editing on colonies revealed editing across most colonies with a predominance of homozygous edits (Figure 3F), as expected by our high editing efficiencies.

**Figure 3.**
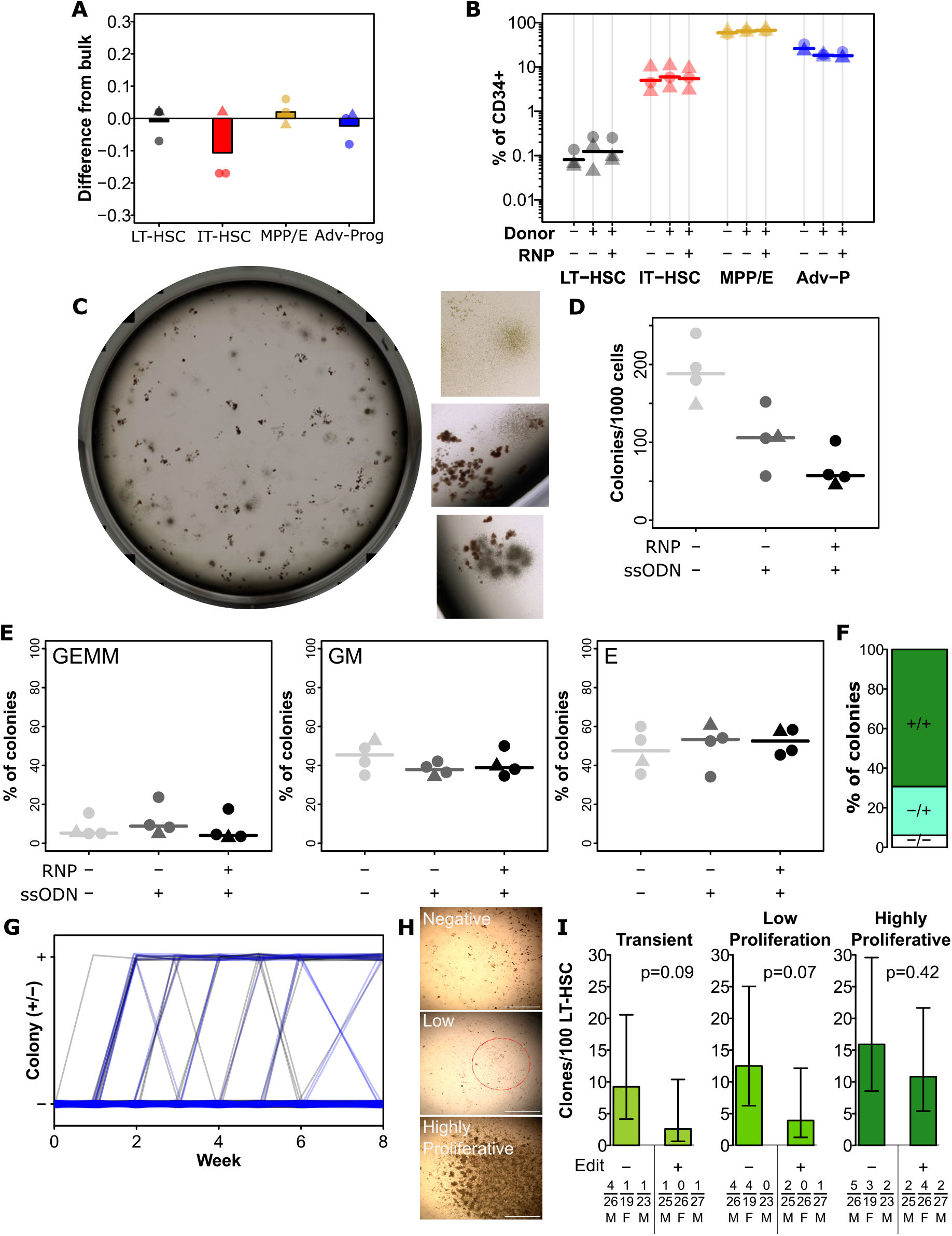
Editing has a minimal impact on HSPC function and hierarchy. **A)** *Integration efficiency is equivalent across phenotypically defined progenitor compartments*. All edits were performed with 0.5 μM AZD7648, 20 fmol p53 siRNA, and 50 pmol of silent mutation ssODN. Values show the difference in precise edit efficiency for each phenotypic subset compared to bulk assessment within that cord. Bars show mean values and points show measurements for individual cords. Male cords are shown as triangles and female as circles. All populations show no significant difference from bulk. **B)** *Progenitor phenotypes are minimally altered across the hierarchy*. The % of CD34+ for each sub-population is shown. Mean values are indicated as lines. A slight but significant decrease was present for late progenitors (CD34+CD45RA+) associated with donor addition (but not different with editing). **C)** *An example image of a well of colonies, and example colonies*. **D)** *Total colonies are decreased by the addition of donors, and further by editing*. Total CFC per 1000 CD34+ cells is shown for each cord. Lines indicate mean values. **E)** *No changes were observed in the frequency of colonies of each type*. As before points are individual cords and lines show mean values. **F)** *Colonies showed a preponderance of homozygous editing*. Mean homozygous, heterozygous edited, and unedited cells are shown from 36 analyzed colonies across 3 independent cords. False-discovery rate (FDR) corrected paired t-test significance values are shown in Table S1. **G)** *No change in the dynamics of colony emergence from single-LT-HSCs in LTC-IC*. The presence or absence of an obvious colony in each well (initially sorted with a single LT-HSC) was scored weekly over the first 6 weeks of the LTC-IC assay, and again at week 8. Clonal outputs are shown as lines with unedited in black and edited in blue. **H)** *Example colonies at 8 weeks*. At 8 weeks, clones were scored as negative (no colony at any point), transient (previous colony without a colony at endpoint), low proliferation (>50 cells, but below confluence), and highly proliferative (confluent). Example images of negative, low proliferation, and highly proliferative clones are shown. Scalebars (white) show 1 mm. The low proliferation colony is circled in red. **I)** *Highly proliferative clones are not lost from the LT-HSC population in the editing process*. The frequency of clones of the indicated types is shown per 100 phenotypic LT-HSC either without editing or following optimal editing. Error bars represent 95% confidence intervals. Frequencies, p-values, and error bars were calculated using Extreme Limiting Dilution Analysis, based on colony numbers measured from 3 independent experiments (each with a different cord donor). Numbers for each clone per donor, the total number of clones analyzed for that donor, and donor sex are indicated below the relevant bar.

In order to assess whether the editing process affected the maintenance of HSC with high self-renewal capacity, we used a clonal version of the long-term culture initiating cell (LTC-IC) assay. We have previously shown that clones with high proliferative potential at 8 weeks in this assay correlate with those HSC with the highest regenerative potential in vivo^18^. Overall, we observed a mixture of negative, transient, low proliferation, and highly proliferative clones across the tested cords and conditions with no difference in the kinetics of clone emergence between edited cells and non-edited controls (Figure 3G,H). Critically, there was no significant difference in the frequency of highly proliferative clones between edited cells and non-edited controls (Figure 3I), suggesting that the editing process does not substantially affect the most regenerative subset of HSC. While there was not a significant difference, there was a trend towards lower frequencies of low proliferation and transient clones in the edited condition (Figure 3I). This is consistent with the observed decrease in CFC frequency (Figure 3D), suggesting that later progenitors may be more affected by editing than the most primitive subsets.

### Mutation zygosity can be tuned based on the ratio of mutant to silent donor

While high-level homozygous editing such as those achieved by our current protocol is desired for therapeutic purposes, for disease modelling applications, heterozygosity is often required for accurate modelling. To enable these applications, we tested whether editing efficiency and zygosity could be tuned by providing a mixture of wild-type and mutant donors. To test this, we used a mix of our AAV with different ratios of silent ssODN (Figure 4A). At 48 hours post editing a bulk sample was harvested to determine overall editing efficiency. This showed a highly significant monotonic decrease in mutant allele frequency with increasing silent donor proportion (p<<0.001, Figure 4B). It should be noted that overall editing efficiencies in these tests were lower than normal due to the requirement of simultaneously editing many conditions. To confirm whether the strategy was indeed tuning the zygosity and not simply the overall efficiency, we sorted single CD34+ cells in 96-well plates and allowed these to grow into a colony prior to harvest and genotyping (Figure 4C). No significant differences in clonogenic efficiency were observed across conditions though there were no large clones (ie. in excess of 100 cells) in the 100% mutant donor condition (Figure 4C). These results suggest that this analysis was not confounded by differential clonogenicity of specific mutational statuses (homozygous WT/silent, heterozygous, homozygous mutant). As expected, we observed a preponderance of homozygous edits with all mutant donor, with a progressive increase in heterozygous or homozygous silent as the proportion of silent donor was increased (Figure 4D). Desired zygosity ratios could thus be predicted based on simple sampling calculations.

**Figure 4.**
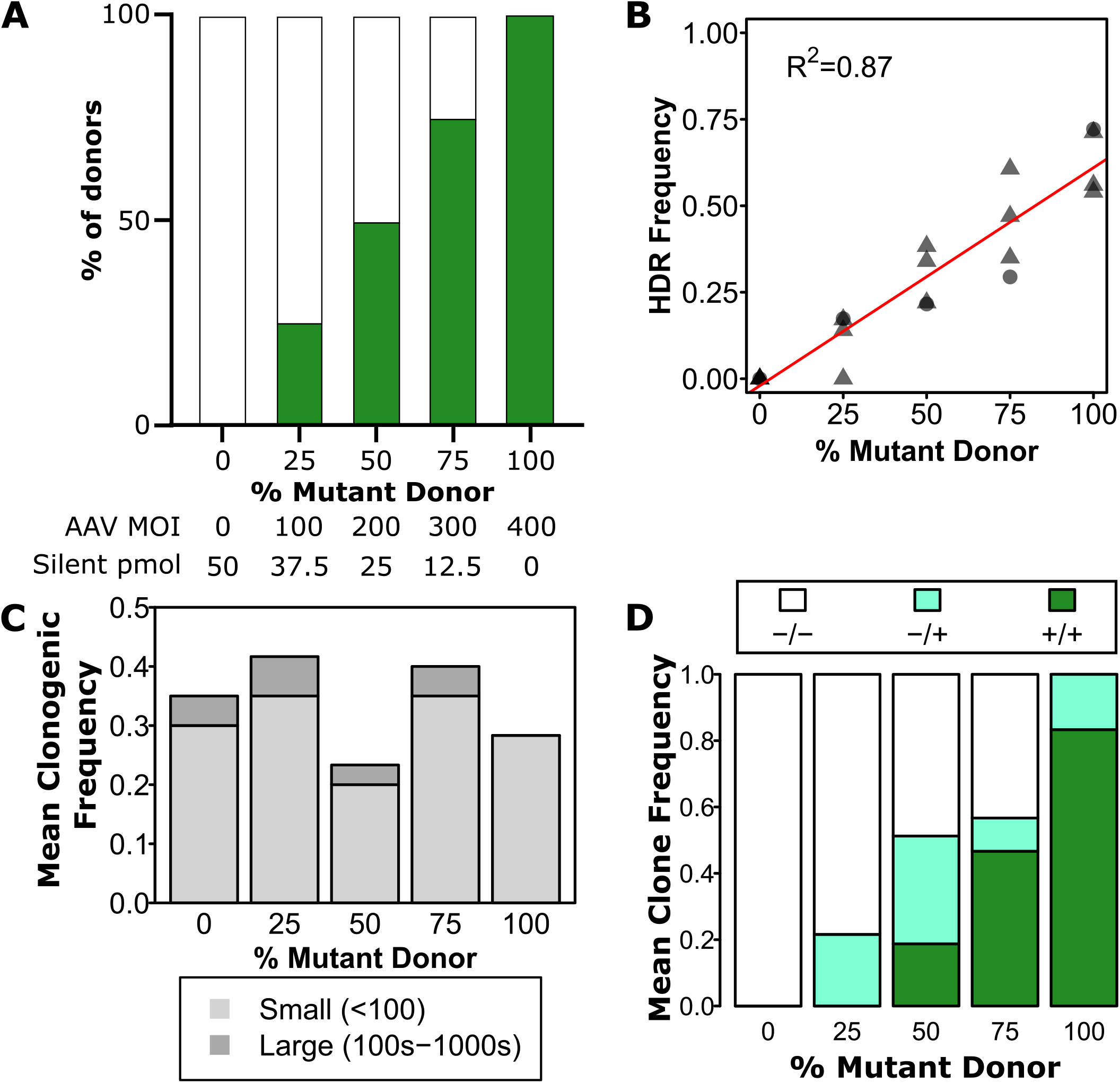
Zygosity can be tuned using a mixture of mutant and silent donors. **A)** *Experimental design*. Green bars represent the proportion of mutant donor. Specific amounts of mutant and silent donor are shown underneath for each condition. All edits were performed with 0.5 μM AZD7648, 20 fmol p53 siRNA, and indicated amounts of each donor. **B)** *Overall mutant integration efficiency varies linearly with the proportion of mutant donor*. Individual cords are shown as points. Male cords are shown as triangles and female as circles. A linear fit is indicated as a red line. The R^2^ is indicated, and overall p value was <<0.001. **C)** *Mean clonogenic frequencies are consistent across donor proportions*. Data are a mean of two independent cords with a total of 30 cells for each cord at each dose analysed except the 0% condition which only had 20 cells per cord. **D)** *Zygosity can be adjusted by inclusion of silent donor*. Mean frequencies of homozygous mutant, heterozygous and homozygous silent donors are shown within all clones with any observed editing. A total of 52 clones with some degree of editing across 2 independent cords were analyzed.

## Discussion

The current study demonstrates a near-perfect efficiency of precise editing in primary human HSPC and identifies the key design and technical considerations needed to achieve it. Starting from existing optimized protocols for AAV-based editing^4^, we tested a variety of dosages, HDR donors, donor designs, and small molecule additives for their ability to improve HDR efficiency while retaining cell viability. Through these tests, we confirmed reports that DNA-PK inhibition was an effective method to improve HDR efficiency^10,11^, and identified AZD7648 to be optimal for this task, consistent with demonstrations of its improved potency and specificity^14^. Combining AZD7648 with optimal HDR donors enabled both AAV and ssODN donors to achieve >90% editing efficiency in HSPC. This is particularly interesting as 53BP1 inhibition was reported to be less effective for ssODN donors^10^. The difference is likely attributable to the design of the ssODN donors themselves, as we also found that additional silent mutations in the spacer sequence were necessary to achieve maximal efficiencies. Given that most AAV donors insert a tag of some description^4,5,8,10^, such a spacer disruption has been built-in to these systems, but not in the competing ssODN. This simple design change facilitates the use of these much cheaper, faster to iterate, and more stable donors. One surprising factor that had a substantial effect on efficiency was the number of conditions simultaneously processed at the time of nucleofection. This likely corresponds to the handling time, and may be analogous to the delivery window for AAV immediately post-nucleofection previously reported^5^.

Beyond simple viability and efficiency, CD34+ cells (HSPCs) are a heterogeneous population made up of a continuous hierarchy of progenitors and stem cells defined by their functional capabilities. To ensure that the editing process was functional throughout the hierarchy, and not specifically toxic to specific levels, we demonstrated that editing did not change the frequency of populations across this hierarchy and that editing efficiency was equivalent throughout, including in the HSC phenotype. These tests made use of phenotypic markers previously shown to be stable and functionally predictive in cultured HSPC^23^. Beyond this, using clonogenic assays for progenitor differentiation, we demonstrated that editing did not alter the distribution of myeloid outputs. These findings together with previous reports demonstrating that editing rates were consistent in pre-transplant measured and post transplantation both with and without DNA-PK inhibition^4,5,9,10^ suggest that the reported editing protocol also edited cells capable of engrafting a mouse for >6 months (functional long-term HSC). This is consistent with the maintenance of highly proliferative LTC-IC we observed. That said, long-term transplants were not performed here. It thus remains a formal possibility that functionally defined LT-HSC may either be edited at lower frequency or functionally compromised, though our clonal LTC-IC data would suggest that this is not the case. We did, however, observe a modest decrease in CFC potential and a non-significant trend towards the same in low proliferation LTC-IC, suggesting that late progenitors may be more sensitive to the editing process than the most primitive. This would be consistent with the greater sensitivity of late progenitors to even growth factor stimulation which detrimentally affects their engraftment ability^27^. It is important to note, however, that our and others’ data suggest that remaining CFC appear to retain normal function^9,10^. Interestingly, much of the toxicity both in direct cell survival and CFC numbers could be attributed to donor delivery, regardless of editing. This suggests that toxicity is likely mediated by innate immune responses to DNA^28^. Interference with these pathways may thus be a fruitful avenue to mitigate toxicity and should be explored in future studies.

Of course, several other factors remain important considerations for specific use-cases. For disease modelling, zygosity can be an important factor, and thus having 100% mutant may not be ideal. To address this issue, we demonstrated that by delivering a combination of silent and mutant donors (of the same strand to prevent donor annealing) one can re-introduce heterozygosity based on the ratio of donors delivered, allowing editing levels and zygosity to be tuned. Another consideration that would be important for any clinical use of edited cells is off-target editing and genomic rearrangements^29^. We did not observe any off-target edits at predicted sites, regardless of whether edits were done in the presence of AZD7648/p53 siRNA or not, suggesting that the protocol presented here did not substantially increase off-target editing frequencies. This was consistent with findings from other studies using 53BP1 inhibition which similarly targets NHEJ, where they observed no increases in off target editing or large-scale genomic rearrangements were observed when donor was present^10^.

During the time that this article was undergoing its publication process, another manuscript came out showing similar effects for the addition of AZD7648 as being able to increase the precise editing efficiency of human HSPCs (as well as T and B cells, and induced pluripotent stem cells)^30^. In their manuscript they also show an increase in efficiency to similar levels using AAV6 donors, and similar but slightly lower inefficiencies that we achieved in ssODN donors^30^. The differences are likely attributable to our use of shorter ssODN donors and our choice of silent spacer disrupting mutations in them. Both of these factors have been previously shown to impact the integration efficiency^31^. We also observed similar viability effects of AAV donor, and lack of negative effects from 0.5 μM AZD7648^30^, though we extend this with the effects of the ssODN donors and ssODN donor length on viability. Again, we similarly show that CFC distributions are not affected by the editing process^30^, though we again extend this with information on the effect of ssODN donors on total CFC frequency. Beyond what was done in the other manuscript, we also show that the editing process and addition of AZD7648 edits evenly across the HSPC hierarchy, and that it does not substantially affect the distribution of progenitor phenotypes and critically does not affect the LTC-ICs with the highest self-renewal potential. Also unique in our manuscript, we demonstrate that for use in mutation modeling/leukemia where zygosity can be an important consideration, it’s possible to tune the zygosity of output cells by adjusting the ratios of mutant to silent donors. Finally, another article was also published while we were in our review process in which they used AZD7648 together with Polθ inhibitors in several non-hematopoietic cell lines to improve editing efficiency^32^. Overall, these and our manuscripts cross-validate the use of AZD7648 for improving precise genome editing in HSPC and beyond (and across more loci, as ours differ completely from theirs^30,32^).

Overall, this study identified a number of critical factors that determine the efficiency of precise editing in primary human HSPCs, and how the combination of these can allow editing at near-perfect efficiency. Moreover, this efficiency could be tuned by the inclusion of competing silent donor. We anticipate that the protocol presented here will enable precise isogenic disease modelling for a variety of monogenic blood diseases directly in primary human HSPCs. The principles and protocols will also permit further improvement in existing approaches for therapeutic editing that are even now in trial.

## Author Contributions

FMCU: Conceptualization, Methodology, Investigation, Writing Original – Original Draft, Writing - Review & Editing, Visualization, Data Curation. JB: Investigation, Writing - Review & Editing. AS: Investigation, Writing - Review & Editing, BL: Methodology, Resources, Writing - Review & Editing. GS: Supervision, Resources, Writing - Review & Editing. HS: Supervision, Conceptualization, Methodology, Writing - Review & Editing. DJHFK: Conceptualization, Methodology, Formal Analysis, Writing Original – Original Draft, Writing - Review & Editing, Visualization, Supervision, Project administration, Funding acquisition

## Acknowledgements

Funds for this work were provided from the IRIC Philanthropic funds from the Marcelle and Jean Coutu foundation, Fonds Innovation du Groupe Canam, the Cancer Research Society (Operating Grants program #944254), and the Terry Fox Research Institute (Terry Fox New Investigator Award #TFRI 1118). D.J.H.F.K. has salary support from FRQS in the form of a Chercheurs-boursiers Junior 1 fellowship (#283502). F.M.C.U. was supported by a Cole Foundation Doctoral Award, a bourse d’excellence du programme de biologie moléculaire from the Université de Montréal and a PhD scholarship from the Institut de Recherche en Immunologie et en Cancérologie.

The authors would like to thank Thomas Sontag and Michel Duval from the Banque de Recherche de Sang de Cordon at CHU Sainte-Justine and HemaQuebec for assistance with cord blood acquisition, the generous individuals who donated their cords for this project, Annie Gosselin and Angélique Bellemare-Pelletier from the IRIC Flow Cytometry facility for technical assistance with sorts, Raphaelle Lambert from the IRIC Genomics facility for technical assistance with Sanger sequencing, the Canadian Neurophotonics Platform Viral Vector Core Facility (RRID:SCR_016477) for assistance with AAV generation. AAV2_HLF-ZE which was used to generate the SRSF2 AAV donor was a gift from Guy Sauvageau (Addgene plasmid # 175034; http://n2t.net/addgene:175034;RRID:Addgene_175034). The LTC-IC feeders were a generous gift from Dr. Connie J. Eaves provided with the assistance of Glenn Edin and Margarita E MacAldaz.

