## Supplementary for "Near-perfect precise on-target editing of human hematopoietic stem and progenitor cells"

**Supplementary Information**

**Key resources table**

| REAGENT or RESOURCE | SOURCE | IDENTIFIER |
|---|---|---|
| Antibodies | | |
| AF647 Mouse Anti-Human CD34 (clone 581) | Cedarlane | 343508 |
| V450 Mouse Anti-Human CD45RA (clone HI100) | BD Biosciences | 560362 |
| PE-CF594 Mouse Anti-Human CD90 (clone 5E10) | BD Biosciences | 562385 |
| FITC anti-human, CD49c (Clone REA360) | Miltenyi | 130-105-364 |
| Bacterial and virus strains | | |
| Custom AAV6 – pssAAV_SRSF2P95H | Canadian Neurophotonics Platform – Viral Vector Core | Custom AAV |
| Biological samples | | |
| Human Cord Blood for CD34+ cells harvest | Héma Québec via St Justine hospital | |
| Chemicals, peptides, and recombinant proteins | | |
| Alt-R® S.p. Cas9 Nuclease V3, 500 µg | IDT | 1081058 |
| T7 Endonuclease I - 250 units | NEB | M0302S |
| Proteinase K, Molecular Biology Grade | NEB | P8107S |
| Alt-R® Cas9 Electroporation Enhancer, 2 nmol | IDT | 1075915 |
| PLATINUM SUPERFI II MASTER MIX | Life Technologies | 12368050 |
| BspEI Enzyme | NEB | R0540S |
| StemSpan™ CC100 | STEMCELL Technologies | 2690 |
| UM 171 | ExcellThera | |
| RS-1 | Cedarlane | 21037-5 |
| Nedisertib (M3814) | Cedarlane | A17055 |
| AZD 7648 | Cedarlane (Cayman) | 28598-1 |
| p53 siRNA id s605 | Thermo | 4390824 |
| DIMETHYL SULFOXIDE (DMSO), Sterile | BioShop | DMS666.100 |
| IL-3, Human (CHO-expressed), 100 ng/ul | Cedarlane (GeneScript) | Z02991-10 |
| SCF, Human (P. pastoris-expressed), 100 ng/ul | Cedarlane (GeneScript) | Z02692-10 |
| EPO 100 ng/ul (~16 IU/uL) | Cedarlane (GeneScript) | Z02975-10 |
| Flt-3L 100 ng/ul | Cedarlane (GeneScript) | Z02926-10 |
| GM-CSF, Human (CHO-expressed), 100 ng/uL | Cedarlane (GeneScript) | Z02983-10 |
| IL-6, Human (CHO-expressed), 100 ng/uL | Cedarlane (GeneScript) | Z03134-50 |

| | | |
|---|---|---|
| G-CSF, Human (CHO-expressed), 100 ng/uL | Cedarlane (GeneScript) | Z02980-10 |
| Hydrocortisone | BioShop | HYD400.5 |
| MyeloCult™ H5100 | STEMCELL Technologies | 05150 |
| CellAdhere™ Type I Collagen, Bovine, Solution | STEMCELL Technologies | 7001 |
| Blunt/TA Ligase Master Mix | NEB | M0367S |
| NEBNext® Quick Ligation Module | NEB | E6056S |
| NEBNext® Ultra™ II End Repair/dA-Tailing Module | NEB | E7546S |
| 1M buffer | Homemade | Homemade |
| Critical commercial assays | | |
| EasySep™ Human CD34 Positive Selection Kit II | STEMCELL Technologies | 17896 |
| MethoCult™ H4034 Optimum | STEMCELL Technologies | 4034 |
| Flongle Sequencing Expansion | Oxford Nanopore | FLO-FLG114 |
| Flongle Flow Cell (R10.4.1) | Oxford Nanopore | FLO-FLG114 |
| Native Barcoding Kit 24 V14 | Oxford Nanopore | SQK-NBD114.24 |
| Software and algorithms | | |
| GelAnalyzer 19.1 | Istvan Lazar Jr. and Istvan Lazar Sr. | www.gelanalyzer.com |
| Synthego Performance Analysis V3 | ICE Analysis | https://www.synthego.com |
| FlowJo™ Software 10.8.1 | BD Life Sciences | https://www.flowjo.com |
| R (version 4.1.2) | R Core Team | https://www.r-project.org/ |
| MinKNOW (version 23.11.3) | Oxford Nanopore | https://community.nanoporetech.com/downloads |
| minimap2 (version 2.26) | Dana-Farber Cancer Institute | https://github.com/lh3/minimap2 |
| samtools (version 1.10) | Genome Research Ltd. | http://www.htslib.org/ |
| bcftools (version 1.10.2) | Genome Research Ltd. | https://samtools.github.io/bcftools/ |
| Biopython (version 1.83) | Biopython | https://biopython.org/ |
| Python (version 3.9.7) | Python | https://www.python.org/ |
| Other | | |
| FBS Canadien | Thermo | 12483020 |
| RPMI1640 | LifeTech | 11875119 |
| StemSpan™ SFEM II | STEMCELL Technologies | 9655 |
| M210B4 expressing human IL-3 and G-CSF | Gift from Connie J Eaves | |
| sl/sl mouse fibroblasts expressin human SCF and IL-3 | Gift from Connie J Eaves | |
| sl/sl mouse fibroblasts expressin human FLT3L | Gift from Connie J Eaves | |

### Oligonucleotides (all from IDT)

| Name | Description | Sequence |
|---|---|---|
| SRSF2_gRNA1 | crRNA SRSF2 | /AlTR1/rCrGrGrCrUrGrUrGrGrUrGrUrGrArGrUrCrCrGrGrGrUrUrUrUrArGrArGrCrUrArUrGrCrU/AlTR2/ |
| pri0077-F | Outer PCR SRSF2 | AGCGATATAAACGGGCGCAG |
| pri0077-R | Outer PCR SRSF2 | TCGCGACCTGGATTTGGATT |
| pri0002-H3 | Inner PCR SRSF2 | CTATGGATGCCATGGACGGG |
| pri0002-H4 | Inner PCR SRSF2 | CAAGCACAGCGGGGTTAATTC |
| pri0261-F | SRSF2 gRNA1 off-target 1 | TCATTGGCAAACAGCAAGCC |
| pri0261-R | SRSF2 gRNA1 off-target 1 | AGAAGTATGTGCCTACGCGG |
| pri0262-F | SRSF2 gRNA1 off-target 2 | GAGAGTCACCGACCATGACG |
| pri0262-R | SRSF2 gRNA1 off-target 2 | TGTAAAACGTGCTGGAGGCT |
| pri0263-F | SRSF2 gRNA1 off-target 3 | CAGAAAGCACAAGCAACGCT |
| pri0263-R | SRSF2 gRNA1 off-target 3 | TCTCTTCCGGACACAAGTGC |
| pri0285 | SRSF2 off-target 2 sequencing | CTCCTTCTTCACGTCTTCCT |
| pri0286 | SRSF2 off-target 3 sequencing | CACCACATCTGGGATCCTCA |
| SF3B1 Cas9 gRNA K700 | crRNA SF3B1 | /AlTR1/rUrGrGrArUrGrArGrCrArGrCrArGrArArArGrUrUrGrUrUrUrArGrArGrCrUrArUrGrCrU/AlTR2/ |
| Alt-R® CRISPR-Cas9 tracrRNA | tracrRNA | Cat #: 1072533 |
| pri0078-F | Outer PCR SF3B1 | GCTGCTGGTCTGGCTACTAT |
| pri0078-R | Outer PCR SF3B1 | ATACTCATTGCTGATTACGTGATTT |
| pri0002-H1 | Inner PCR SF3B1 | TGGGCTACTGATTTGGGGAG |
| pri0002-H2 | Inner PCR SF3B1 | CTGTGTTGGCGGATACCCTT |

**Donor DNA sequences**

**Silent mutation**

**Mutation**

Synthetic Intron/Inserted Sequence

Guide RNA complementary sequence

SRSF2 Silent ssODN:

tggacggccgcgagctgcgggtgcaaatggcgcgctacggccgcccTccAgaTtcacaccacagccgccggggaccgccaccccgcaggt

SRSF2 P95H ssODN:

tggacggccgcgagctgcgggtgcaaatggcgcgctacggccgccATccggactcacaccacagccgccggggaccgccaccccgcaggt

SRSF2 long ssODN donor

ttcacgacaagcgcgacgctgaggacgctatggatgccatggacggggccgtgctggacggccgcgagctgcgggtgcaaatggcgcgctacggcc gccATccggactcacaccacagccgccggggaccgccaccccgcaggtacgggggcggtggctacggacgccggagccgcaggtaaacggggctg aggggaccg

SRSF2 P95H ssODN with additional silent mutations:

tggacggccgcgagctgcgggtgcaaatggcgcgctacggccgccATccAgaTtcacaccacagccgccggggaccgccaccccgcaggt

SRSF2 P95H AAV donor sequence:

cctgcaggcagctgcgcgctcgctcgctcactgaggccgcccgggcgtcgggcgacctttggtcgcccggcctcagtgagcgagcgagcgcgcagag agggagtggccaactccatcactaggggttcctgcggcctctagACCGGCGTCCGTGCTGTTCTGCGGCaaggcctttcccagtgtcccca cgcggaaggcaactgcctgagaggcgcggcgtcgcaccgcccagagctgaggaagccggcgccagttcgcggggctccgggccgccactcagagct atgagctacggccgccccccctcccgatgtggagggtatgacctccctcaaggtggacaacctgacctaccgcacctcgcccgacacgctgaggcgcgt cttcgagaagtacgggcgcgtcggcgacgtgtacatcccgcgggaccgctacaccaaggagtcccgcggcttcgccttcgttcgctttcacgacaagc gcgacgctgaggacgctatggatgccatggacggggccgtgctggacggccgcgagctgcgggtgcaaatggcgcgctacggccgccAcccggtA AGTgAAAAAagcatagctctaaaacTGCTTCGCTACTGCATCGGCCGGGAATCGAACCCGGGCCGCCCGCGTGGCAGG CGAGCATTCTACCACTGAACCACCGATGCTACTAACTCGAGagTTCTTTCTTTCTTTCACAGgactcacaccacagccgccg gggaccgccaccccgcaggtacgggggcggtggctacggacgccggagccgcaggtaaacggggctgaggggaccgcgggaggcggggcggggc gcgcgggaggcccgggcgacctcacaaaggtccgcggcgaagcacgtggtgcgggcccggacggggcgggggtgcacgccgcgtctcgcgaccct ccggccaccccgcgagcttccgccgtctgcgacccgggagtggccggggtgtgggcggcgcggggcggaggaccccgcctcgcgactggggaaatg gcgtctggcggcgagataatggcggcctgggcgggagcgcgcgggcggggccggccccgctgcctggaattaaccccgctgtgcttgctcgtcccgcc cgcagccctaggcggcgtcgccgcagccgatcccggagtcggagtcgttccaggtctcgcagccgatctcgctacagCCGCTCGAAGTCTCGG TCCCGCTAGTgcggccgcaggaacccctagtgatggagttggccactccctctctgcgcgctcgctcgctcactgaggccgggcgaccaaaggtc gcccgacgcccgggctttgcccgggcggcctcagtgagcgagcgagcgcgca

SF3B1 K700E AAV donor sequence

GAGCTTTTGCTGTTGTAGCCTCtgccctgggcattccttctttattgcccttcttaaaagctgtgtgcaaaagcaagaagtcctggcaagcgag acacactggtattaagattgtacaacagatagctattcttatgggctgtgccatcttgccacatcttagaagtttagttgaaatcattgaacatggtaagt tgtaatgtaactttgtctttttttttttttctttaggagacagggtcttactatgttgcccagactggactcaaacctttgggctcaagtgatcctcctgctca gcctccttagtagttgggactagaggtacacacacagcctgtccatgtttaataggacagctgtcctaaaattTGGGCTACTGATTTGGGGAG

ataaatggaaaggcatagctctacaaactatagattttatgatgggtttgttatattatctgctgacaggctatggttcatgttttgcttttacctaattttg
tttaatgtgaacatattctgcagtttggctgaatagttgatatattgagagaatctggatgatattgtgtaacttaggtaatgttggAAAAAagcatag
ctctaaaacTGCTTCGCTACTGCATCGGCCGGGAATCGAACCCGGGCCGCCCGCGTGGCAGGCGAGCATTCTACCAC
TGAACCACCGATGCggcatagttaaaacctgtgtttg**C**ttttgtaggtcttgt**T**gatgagca**A**cag**G**aagttcggaccatcagtgctttggccat
tgctgccttggctgaagcagcaactccttatggtatcgaatcttttgattctgtgttaaagcctttatggAAGGGTATCCGCCAACACAGagga
aaggtaaatccaccaattaccttttgatttatcttcattaaagttaaggcgacataaatctaaattactaaagtacatatattttttatttaaaaataggg
tttggctgctttcttgaaggctattgggtatcttattcctcttatggatgcagaatatgccaactactatactagagaagtgatgttaatccttattcgaga
attccagtctcctgatgaggaaatgaaaaaaattgtgctgaaggtaattattccagatttgttaatgtaaaCTGGATATGTTTCATGGTTCTA
ACATAGT

### Supplementary Tables:

**Table S1. Significance testing.** Figure and panel are indicated along with each pair-wise test. Where relevant the colony type or population for a given test is indicated.

| Figure | Panel | Condition 1 | Condition 2 | Pvalue | FDR | Colony type / Population |
|---|---|---|---|---|---|---|
| 1 | c | SRSF2_AAV_yes_200 | SRSF2_AAV_yes_400 | 0.023277153 | 0.023277153 | NA |
| 1 | c | SRSF2_AAV_yes_400 | SRSF2_AAV_yes_800 | 0.012716956 | 0.023277153 | NA |
| 1 | d | SRSF2_ctrl_no_ 0 | SRSF2_AAV_no_200 | 0.21963085 | 0.241593935 | NA |
| 1 | d | SRSF2_ctrl_no_ 0 | SRSF2_AAV_yes_200 | 0.158741852 | 0.194017819 | NA |
| 1 | d | SRSF2_ctrl_no_ 0 | SRSF2_AAV_no_400 | 0.093385253 | 0.171206296 | NA |
| 1 | d | SRSF2_ctrl_no_ 0 | SRSF2_AAV_yes_400 | 0.068232666 | 0.150111866 | NA |
| 1 | d | SRSF2_ctrl_no_ 0 | SRSF2_AAV_no_800 | 0.002150054 | 0.023650592 | NA |
| 1 | d | SRSF2_ctrl_no_ 0 | SRSF2_AAV_yes_800 | 0.063494273 | 0.150111866 | NA |
| 1 | d | SRSF2_AAV_no_200 | SRSF2_AAV_yes_200 | 0.139663229 | 0.194017819 | NA |
| 1 | d | SRSF2_AAV_no_400 | SRSF2_AAV_yes_400 | 0.040511155 | 0.150111866 | NA |
| 1 | d | SRSF2_AAV_no_800 | SRSF2_AAV_yes_800 | 0.335216014 | 0.335216014 | NA |
| 1 | d | SRSF2_AAV_yes_200 | SRSF2_AAV_yes_400 | 0.152759319 | 0.194017819 | NA |
| 1 | d | SRSF2_AAV_yes_400 | SRSF2_AAV_yes_800 | 0.054599713 | 0.150111866 | NA |
| 1 | e | SRSF2_short_yes_1.00 | SRSF2_short_yes_2.50 | 0.022987461 | 0.045974921 | NA |
| 1 | e | SRSF2_short_yes_2.50 | SRSF2_short_yes_5.00 | 0.12410038 | 0.12410038 | NA |
| 1 | f | SRSF2_ctrl_no_0.00 | SRSF2_short_no_1.00 | 0.083705834 | 0.167411668 | NA |
| 1 | f | SRSF2_ctrl_no_0.00 | SRSF2_short_yes_1.00 | 0.14477643 | 0.183821244 | NA |
| 1 | f | SRSF2_ctrl_no_0.00 | SRSF2_short_no_2.50 | 0.116017299 | 0.17406597 | NA |
| 1 | f | SRSF2_ctrl_no_0.00 | SRSF2_short_yes_2.50 | 0.11604398 | 0.17406597 | NA |
| 1 | f | SRSF2_ctrl_no_0.00 | SRSF2_short_no_5.00 | 0.08124549 | 0.167411668 | NA |
| 1 | f | SRSF2_ctrl_no_0.00 | SRSF2_short_yes_5.00 | 0.063769856 | 0.167411668 | NA |
| 1 | f | SRSF2_ctrl_no_0.00 | SRSF2_long_no_0.25 | 0.057681145 | 0.167411668 | NA |
| 1 | f | SRSF2_ctrl_no_0.00 | SRSF2_long_yes_0.25 | 0.11508911 | 0.17406597 | NA |
| 1 | f | SRSF2_ctrl_no_0.00 | SRSF2_long_no_0.50 | 0.054620275 | 0.167411668 | NA |
| 1 | f | SRSF2_ctrl_no_0.00 | SRSF2_long_yes_0.50 | 0.070837822 | 0.167411668 | NA |
| 1 | f | SRSF2_ctrl_no_0.00 | SRSF2_long_no_1.00 | 0.036009341 | 0.167411668 | NA |
| 1 | f | SRSF2_ctrl_no_0.00 | SRSF2_long_yes_1.00 | 0.041960004 | 0.167411668 | NA |
| 1 | f | SRSF2_short_no_1.00 | SRSF2_short_yes_1.00 | 0.580156436 | 0.580156436 | NA |
| 1 | f | SRSF2_short_no_2.50 | SRSF2_short_yes_2.50 | 0.163396662 | 0.183821244 | NA |
| 1 | f | SRSF2_short_no_5.00 | SRSF2_short_yes_5.00 | 0.041119811 | 0.167411668 | NA |
| 1 | f | SRSF2_long_no_0.25 | SRSF2_long_yes_0.25 | 0.405690593 | 0.429554745 | NA |
| 1 | f | SRSF2_long_no_0.50 | SRSF2_long_yes_0.50 | 0.159350061 | 0.183821244 | NA |
| 1 | f | SRSF2_long_no_1.00 | SRSF2_long_yes_1.00 | 0.144754023 | 0.183821244 | NA |
| 2 | a | w/o mol | 0.5µM M3814 | 0.028352285 | 0.045363655 | NA |
| 2 | a | w/o mol | 0.5µM AZD7648 | 0.00649234 | 0.014454949 | NA |
| 2 | a | w/o mol | 5µM M3814 | 0.007227475 | 0.014454949 | NA |
| 2 | a | w/o mol | 5µM AZD7648 | 0.003966845 | 0.014454949 | NA |
| 2 | a | 0.5µM M3814 | 0.5µM AZD7648 | 0.001575337 | 0.012602695 | NA |
| 2 | a | 0.5µM AZD7648 | 5µM AZD7648 | 0.139940447 | 0.159931939 | NA |
| 2 | a | 0.5µM AZD7648 | 5µM M3814 | 0.235621749 | 0.235621749 | NA |
| 2 | a | 5µM AZD7648 | 5µM M3814 | 0.133223454 | 0.159931939 | NA |
| 2 | b | Donor only | w/o mol | 0.051358419 | 0.085597365 | NA |
| 2 | b | Donor only | 0.5µM M3814 | 0.035638736 | 0.071277472 | NA |
| 2 | b | Donor only | 0.5µM AZD7648 | 0.150783926 | 0.172700095 | NA |
| 2 | b | Donor only | 5µM M3814 | 0.002710079 | 0.013550395 | NA |
| 2 | b | Donor only | 5µM AZD7648 | 0.033072608 | 0.071277472 | NA |
| 2 | b | 0.5µM M3814 | 0.5µM AZD7648 | 0.00466545 | 0.0155515 | NA |
| 2 | b | 0.5µM M3814 | 5µM AZD7648 | 0.577112708 | 0.577112708 | NA |
| 2 | b | 0.5µM M3814 | 5µM M3814 | 0.155430085 | 0.172700095 | NA |
| 2 | b | 5µM M3814 | 5µM AZD7648 | 0.147738595 | 0.172700095 | NA |
| 2 | b | 0.5µM AZD7648 | 5µM AZD7648 | 0.001816811 | 0.013550395 | NA |
| 2 | c | p53 only_SRSF2_AAV_mutant | p53+RS-1_SRSF2_AAV_mutant | 0.482871408 | 0.482871408 | NA |

| | | | | | | |
|---|---|---|---|---|---|---|
| 2 | c | p53 only_SRSF2_AAV_mutant | p53+AZD_SRSF2_AAV_mutant | 0.06039884 | 0.181196519 | NA |
| 2 | c | p53 only_SRSF2_AAV_mutant | p53+AZD+RS-1_SRSF2_AAV_mutant | 0.242830736 | 0.364246103 | NA |
| 2 | d | Donor only_SRSF2_AAV_ | p53 only_SRSF2_AAV_mutant | 0.740322668 | 0.740322668 | NA |
| 2 | d | Donor only_SRSF2_AAV_ | p53+RS-1_SRSF2_AAV_mutant | 0.156261873 | 0.25411052 | NA |
| 2 | d | Donor only_SRSF2_AAV_ | p53+AZD_SRSF2_AAV_mutant | 0.441147747 | 0.514672372 | NA |
| 2 | d | Donor only_SRSF2_AAV_ | p53+AZD+RS-1_SRSF2_AAV_mutant | 0.069699742 | 0.162632731 | NA |
| 2 | d | p53 only_SRSF2_AAV_mutant | p53+RS-1_SRSF2_AAV_mutant | 0.017514729 | 0.061301553 | NA |
| 2 | d | p53 only_SRSF2_AAV_mutant | p53+AZD_SRSF2_AAV_mutant | 0.181507515 | 0.25411052 | NA |
| 2 | d | p53+AZD_SRSF2_AAV_mutant | p53+AZD+RS-1_SRSF2_AAV_mutant | 0.010588436 | 0.061301553 | NA |
| 2 | g | p53+AZDlow_SRSF2_AAV_mutant | p53+AZDlow_SRSF2_short oligo_mutant | 1.48E-06 | 5.94E-06 | NA |
| 2 | g | p53+AZDlow_SRSF2_AAV_mutant | p53+AZDlow_SRSF2_short oligo_mutantNT | 0.02428539 | 0.03238052 | NA |
| 2 | g | p53+AZDlow_SRSF2_AAV_mutant | p53+AZDlow_SRSF2_short oligo_silent | 0.011248813 | 0.022497626 | NA |
| 2 | g | p53+AZDlow_SRSF2_short oligo_mutantNT | p53+AZDlow_SRSF2_short oligo_silent | 0.058467188 | 0.058467188 | NA |
| 2 | h | SRSF2OT1 Unedited | SRSF2OT1 Edited | 0.23025937 | 0.46051874 | NA |
| 2 | h | SRSF2OT1 Unedited | SRSF2OT1 Edited + AZD7648 | 0.054316581 | 0.325899489 | NA |
| 2 | h | SRSF2OT2 Unedited | SRSF2OT2 Edited | 0.189207494 | 0.46051874 | NA |
| 2 | h | SRSF2OT2 Unedited | SRSF2OT2 Edited + AZD7648 | 0.769245772 | 0.769245772 | NA |
| 2 | h | SRSF2OT3 Unedited | SRSF2OT3 Edited | 0.459381957 | 0.551258348 | NA |
| 2 | h | SRSF2OT3 Unedited | SRSF2OT3 Edited + AZD7648 | 0.449937162 | 0.551258348 | NA |
| 3 | a | LT-HSC | | 0.770584266 | 0.770584266 | NA |
| 3 | a | IT-HSCs | | 0.234177964 | 0.664959451 | NA |
| 3 | a | MPP | | 0.477767032 | 0.664959451 | NA |
| 3 | a | Prog | | 0.498719588 | 0.664959451 | NA |
| 3 | b | Ctrl elec | donor only | 0.348617206 | 0.398419665 | LT-HSC |
| 3 | b | Ctrl elec | donor only | 0.107055326 | 0.171288522 | IT-HSC |
| 3 | b | Ctrl elec | donor only | 0.176332651 | 0.235110201 | MPP |
| 3 | b | Ctrl elec | donor only | 0.088854072 | 0.171288522 | Prog |
| 3 | b | Ctrl elec | Silent | 0.084251875 | 0.171288522 | LT-HSC |
| 3 | b | Ctrl elec | Silent | 0.47964894 | 0.47964894 | IT-HSC |
| 3 | b | Ctrl elec | Silent | 0.014739174 | 0.058956698 | MPP |
| 3 | b | Ctrl elec | Silent | 0.000888205 | 0.007105639 | Prog |
| 3 | d | Ctrl elec | sil w/o RNP | 0.062919995 | 0.067769255 | NA |
| 3 | d | Ctrl elec | sil w RNP | 0.012072095 | 0.036216285 | NA |
| 3 | d | sil w/o RNP | sil w RNP | 0.067769255 | 0.067769255 | NA |
| 3 | e | Ctrl elec | sil w/o RNP | 0.118180694 | 0.531813123 | GEMM |
| 3 | e | Ctrl elec | sil w RNP | 0.588118599 | 0.731982001 | GEMM |
| 3 | e | sil w/o RNP | sil w RNP | 0.013453235 | 0.121079113 | GEMM |
| 3 | e | Ctrl elec | sil w/o RNP | 0.251099079 | 0.731982001 | GM |
| 3 | e | Ctrl elec | sil w RNP | 0.530924473 | 0.731982001 | GM |
| 3 | e | sil w/o RNP | sil w RNP | 0.608350943 | 0.731982001 | GM |
| 3 | e | Ctrl elec | sil w/o RNP | 0.659028761 | 0.731982001 | E |
| 3 | e | Ctrl elec | sil w RNP | 0.465740918 | 0.731982001 | E |
| 3 | e | sil w/o RNP | sil w RNP | 0.731982001 | 0.731982001 | E |
| S1 | a | 30.5_- | 30.5_+ | 0.005011906 | 0.020047625 | NA |
| S1 | a | 30.5_- | 61.0_- | 0.046130418 | 0.084414376 | NA |
| S1 | a | 30.5_- | 61.0_+ | 0.063310782 | 0.084414376 | NA |
| S1 | a | 61.0_- | 61.0_+ | 0.482327699 | 0.482327699 | NA |
| S1 | b | 30.5_- | 30.5_+ | 0.009085768 | 0.012114358 | NA |
| S1 | b | 30.5_- | 61.0_- | 0.390896739 | 0.390896739 | NA |
| S1 | b | 30.5_- | 61.0_+ | 0.000985007 | 0.00328815 | NA |
| S1 | b | 61.0_- | 61.0_+ | 0.001644075 | 0.00328815 | NA |
| S1 | a | 30.5_- | 30.5_+ | 0.113475776 | 0.151301035 | NA |
| S1 | a | 30.5_- | 61.0_- | 0.468501111 | 0.468501111 | NA |
| S1 | a | 30.5_- | 61.0_+ | 0.04124467 | 0.146545815 | NA |
| S1 | a | 61.0_- | 61.0_+ | 0.073272907 | 0.146545815 | NA |
| S1 | d | 30.5_- | 30.5_+ | 0.001208839 | 0.002417678 | NA |
| S1 | d | 30.5_- | 61.0_- | 0.472774519 | 0.472774519 | NA |
| S1 | d | 30.5_- | 61.0_+ | 3.87E-05 | 0.000154923 | NA |
| S1 | d | 61.0_- | 61.0_+ | 0.003437052 | 0.004582736 | NA |
| S3 | a | p53 only_SF3B1_AAV_mutant | p53+RS-1_SF3B1_AAV_mutant | 0.893320553 | 0.893320553 | NA |
| S3 | a | p53 only_SF3B1_AAV_mutant | p53+AZD_SF3B1_AAV_mutant | 2.13E-06 | 8.53E-06 | NA |
| S3 | a | p53 only_SF3B1_AAV_mutant | p53+AZD+RS-1_SF3B1_AAV_mutant | 0.002997226 | 0.005994452 | NA |

| | | | | | | |
|---|---|---|---|---|---|---|
| S3 | a | p53+AZD_SF3B1_AAV_mutant | p53+AZD+RS-1_SF3B1_AAV_mutant | 0.007684271 | 0.010245695 | NA |
| S3 | b | p53 only_SF3B1_AAV_mutant | p53+RS-1_SF3B1_AAV_mutant | 0.058629221 | 0.100709321 | NA |
| S3 | b | p53 only_SF3B1_AAV_mutant | p53+AZD_SF3B1_AAV_mutant | 0.76158459 | 0.76158459 | NA |
| S3 | b | p53 only_SF3B1_AAV_mutant | p53+AZD+RS-1_SF3B1_AAV_mutant | 0.049168908 | 0.100709321 | NA |
| S3 | b | p53+AZD_SF3B1_AAV_mutant | p53+AZD+RS-1_SF3B1_AAV_mutant | 0.075531991 | 0.100709321 | NA |

**Supplementary Figure Legends:**

**Figure S1. RNP efficiency determination.** Cells were edited with indicated amounts of Cas9 RNP and 1x IDT electroporation enhancer where indicated. Cutting efficiencies as measured by T7E1 are shown in (**A**) for SRSF2 and (**C**) for SF3B1, and number of viable cells in (**B**) for SRSF2 and (**D**) for SF3B1. False-discovery rate (FDR) corrected unpaired t-test significance values are shown in Table S1.

**Figure S2: Assays for the detection of HDR integration. A)** *Nested PCR is required to avoid non-specific amplification*. The upper gel shows that even in the absence of RNP, single primer pairs amplify AAV from the donor. The lower gel demonstrates that with nested PCR this is no longer the case. **B)** *Enzymatic digestion can detect editing down to a minority ~1%.* Known proportions of mutant and wild-type PCR product were mixed, digested with BspEI, and the gels quantified. An example gel is shown on the right, and the quantifications from two independent replicates shown on the left. A perfect measure is shown as a blue line and the linear fit in red. **C)** *Sanger sequencing-based detection of editing*. An example chromatogram from an unedited control, a silent edited sample, and a P95H with spacer-breaking silent mutations edited sample are shown. Locations of mutant bases are indicated with a red box for the location of the P95H mutation, and blue boxes for each of the silent mutations. Editing efficiencies calculated by ICE are indicated for each sequence.

**Figure S3. The addition of AZD7648 also improves editing efficiency at the SF3B1 locus. A)** *HDR efficiency for SF3B1 K700E with combinations of AZD7648, p53 siRNA and RS-1*. Cells were edited with 30.5 pmol RNP (or not as indicated) with 400 MOI of AAV donor in the presence of the indicated additives. AZD7648 was used at 5 µM, p53 siRNA at 20 fmol, and RS-1 at 15 µM. **B)** *Viable cell numbers with additive combinations*. Hemocytometer counts at the time of harvest are shown for each sample from (**A**). False-discovery rate (FDR) corrected unpaired t-test significance values are shown in Table S1.

**Figure S4. Example Sanger sequencing traces for the top 3 predicted off-target sites of the SRSF2 gRNA**. Chromosome number and location displayed are indicated for each site. Dotted lines indicate the expected recognition site based on predictions from Benchling. Off-target site 2 is displayed from the reverse strand. All traces were analyzed using ICE and had an estimated cutting of 0% with $R^2$ values between 0.99 and 1. One example chromatogram for each site is shown, however, 3 independent donors were analyzed for each. Trace views were exported using Chromas.

**Figure S5. Full-length nanopore sequencing traces for the top 3 predicted off-target sites of the SRSF2 gRNA. A)** *Frequency of each base type across off-target amplicons*. Chromosome number and location are displayed above each column of samples with the overall amplicon first and the specific predicted target location in brackets. The frequency of reads at a given base that were either reference allele, a substitution (from reference), an insertion, or a deletion are shown for each base across the amplicon. Only reads with a Phred score of at least 16 at a given base were included. Results are shown for unmanipulated control, standard editing (ie, no AZD7648), and edited with our optimal protocol (including AZD7648) are shown for each of 3 individual cord donors (Samples 1-3). Of these, 2 were male (Samples 1 & 2) and 1 female (Sample 3). **B)** *Read depth Per Individual across each amplicon*. The total read depth of accepted bases (ie. those with a Phred score of at least 16) is shown for each sample and region. Target regions are highlighted in grey.

**Figure S6: Example gating hierarchy.** FMO stands for full-minus-one, where the indicated antibody was left out of the panel.

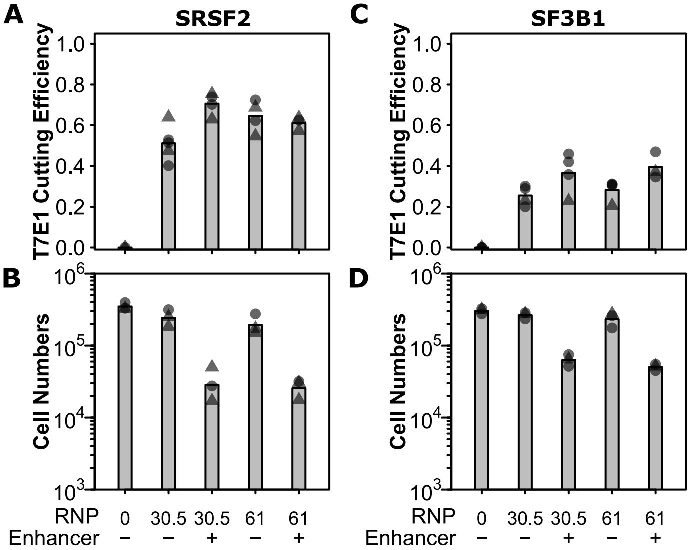

**Figure S1. RNP efficiency determination.** Cells were edited with indicated amounts of Cas9 RNP and 1x IDT electroporation enhancer where indicated. Cutting efficiencies as measured by T7E1 are shown in (**A**) for SRSF2 and (**C**) for SF3B1, and number of viable cells in (**B**) for SRSF2 and (**D**) for SF3B1. False-discovery rate (FDR) corrected unpaired t-test significance values are shown in Table S1.

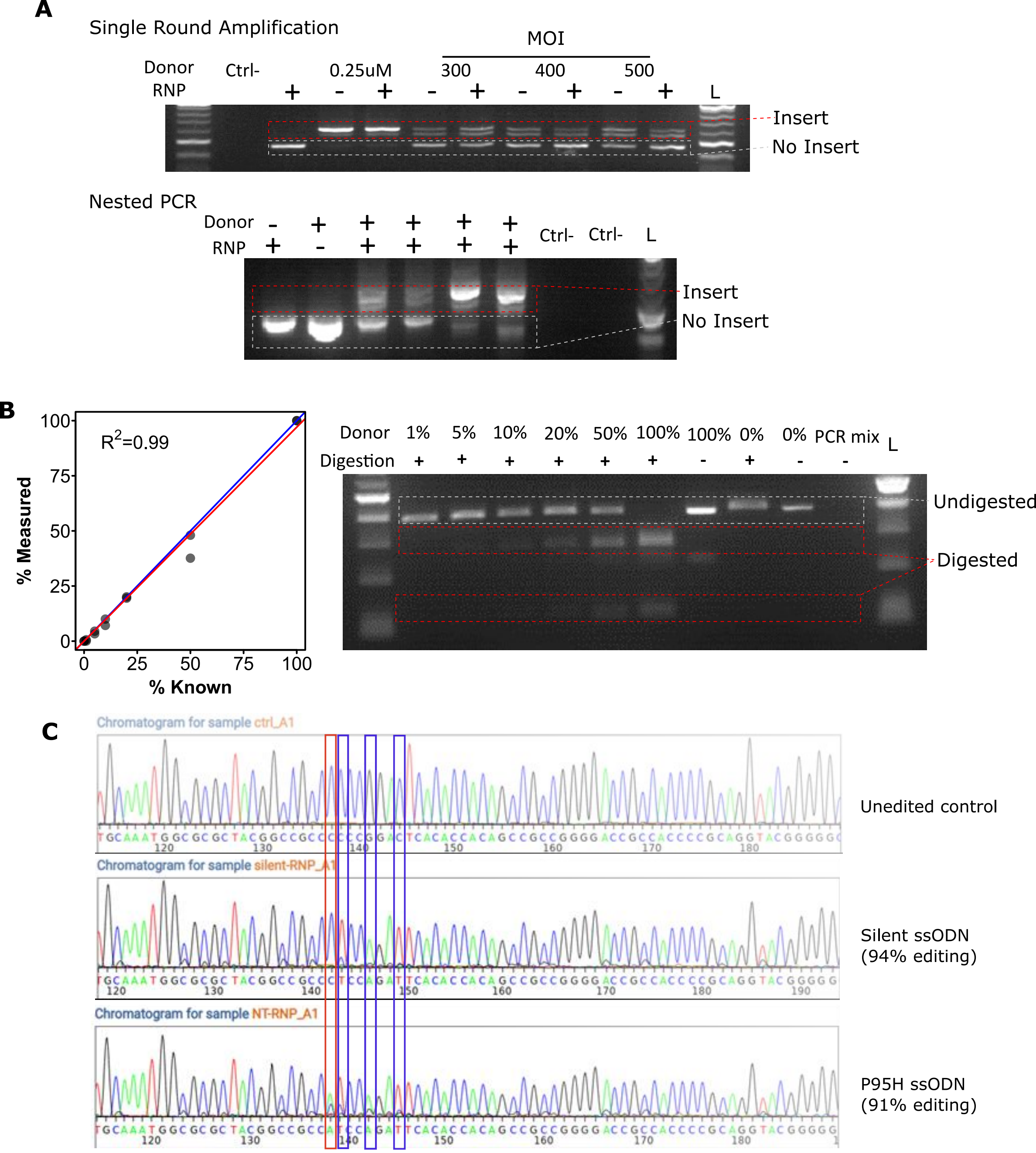

**Figure S2: Assays for the detection of HDR integration. A)** *Nested PCR is required to avoid non-specific amplification*. The upper gel shows that even in the absence of RNP, single primer pairs amplify AAV from the donor. The lower gel demonstrates that with nested PCR this is no longer the case. **B)** *Enzymatic digestion can detect editing down to a minority ~1%*. Known proportions of mutant and wild-type PCR product were mixed, digested with BspEI, and the gels quantified. An example gel is shown on the right, and the quantifications from two independent replicates shown on the left. A perfect measure is shown as a blue line and the linear fit in red. **C)** *Sanger sequencing-based detection of editing*. An example chromatogram from an unedited control, a silent edited sample, and a P95H with spacer-breaking silent mutations edited sample are shown. Locations of mutant bases are indicated with a red box for the location of the P95H mutation, and blue boxes for each of the silent mutations. Editing efficiencies calculated by ICE are indicated for each sequence.

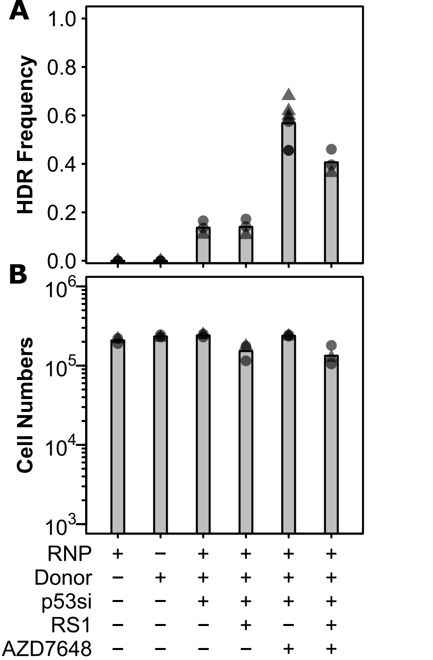

**Figure S3. The addition of AZD7648 also improves editing efficiency at the SF3B1 locus. A)** *HDR efficiency for SF3B1 K700E with combinations of AZD7648, p53 siRNA and RS-1*. Cells were edited with 30.5 pmol RNP (or not as indicated) with 400 MOI of AAV donor in the presence of the indicated additives. AZD7648 was used at 5 μM, p53 siRNA at 20 fmol, and RS-1 at 15 μM. **B)** *Viable cell numbers with additive combinations*. Hemocytometer counts at the time of harvest are shown for each sample from (**A**). False-discovery rate (FDR) corrected unpaired t-test significance values are shown in Table S1.

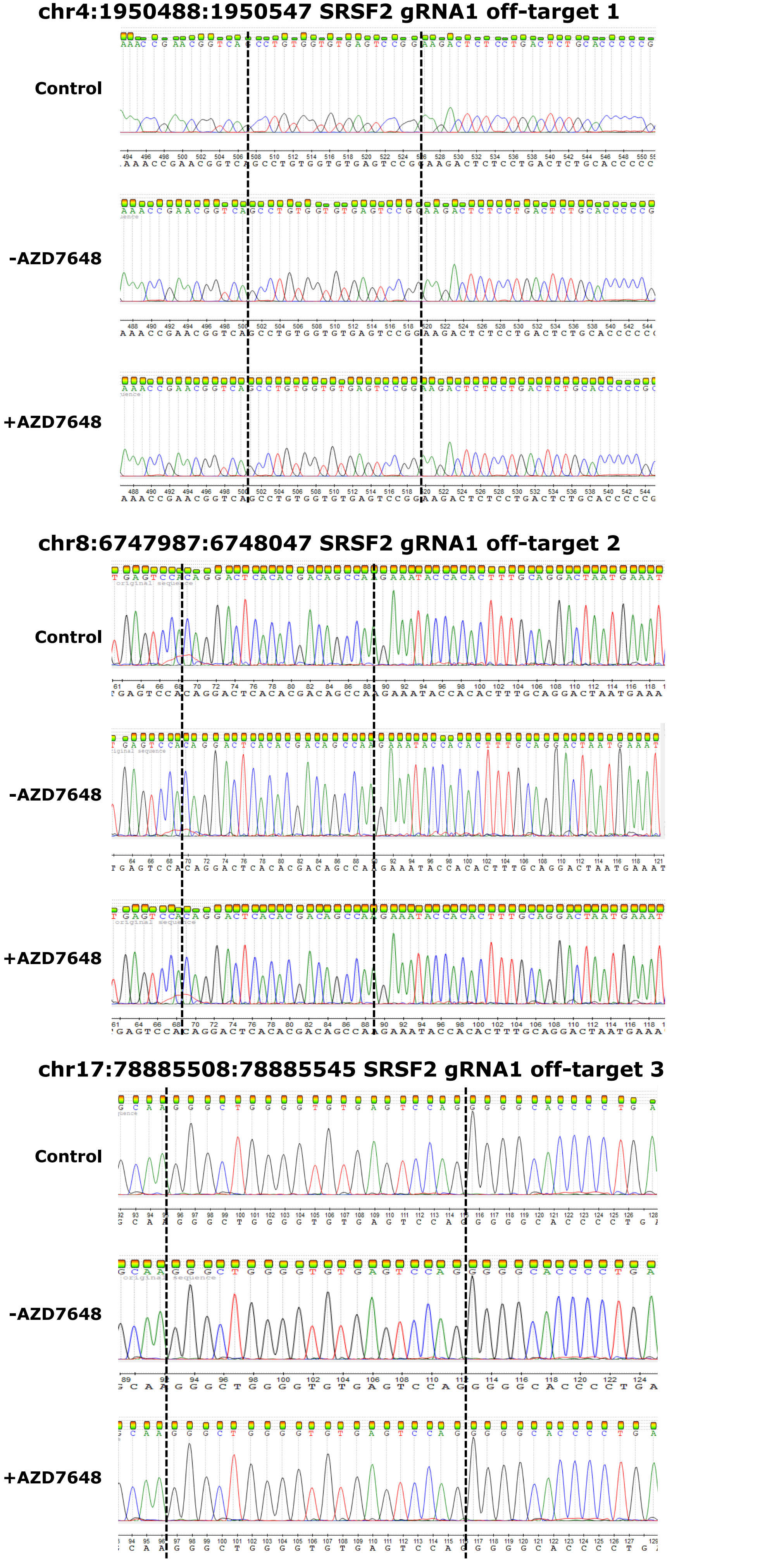

**Figure S4. Example Sanger sequencing traces for the top 3 predicted off-target sites of the SRSF2 gRNA.** Chromosome number and location displayed are indicated for each site. Dotted lines indicate the expected recognition site based on predictions from Benchling. Off-target site 2 is displayed from the reverse strand. All traces were analyzed using ICE and had an estimated cutting of 0% with R2 values between 0.99 and 1. One example chromatogram for each site is shown, however, 3 independent donors were analyzed for each. Trace views were exported using Chromas.

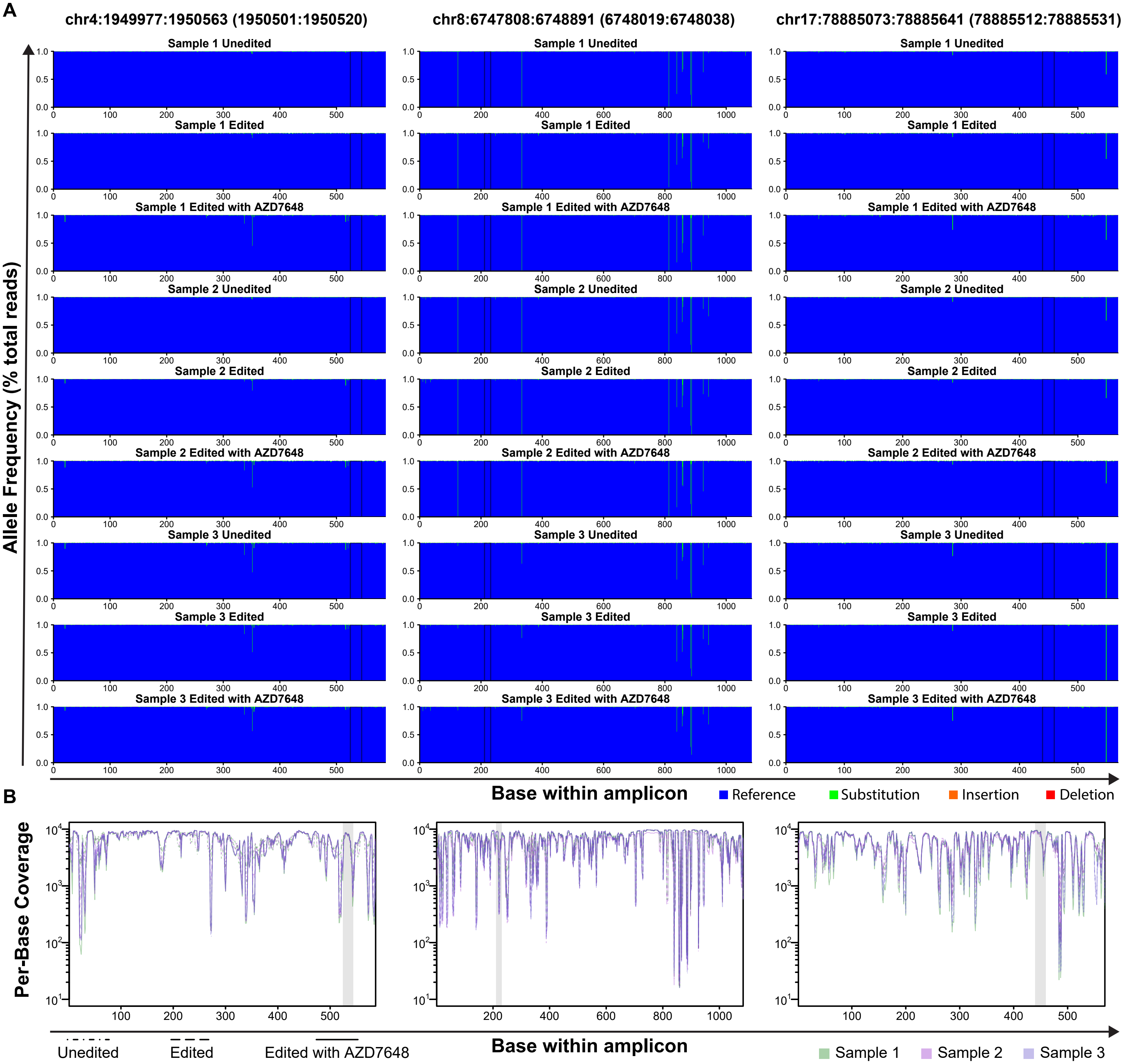

**Figure S5. Full-length nanopore sequencing traces for the top 3 predicted off-target sites of the SRSF2 gRNA. A)** *Frequency of each base type across off-target amplicons.* Chromosome number and location are displayed above each column of samples with the overall amplicon first and the specific predicted target location in brackets. The frequency of reads at a given base that were either reference allele, a substitution (from reference), an insertion, or a deletion are shown for each base across the amplicon. Only reads with a Phred score of at least 16 at a given base were included. Results are shown for unmanipulated control, standard editing (ie, no AZD7648), and edited with our optimal protocol (including AZD7648) are shown for each of 3 individual cord donors (Samples 1-3). Of these, 2 were male (Samples 1 & 2) and 1 female (Sample 3). **B)** *Read depth per individual across each amplicon.* The total read depth of accepted bases (ie. those with a Phred score of at least 16) is shown for each sample and region. Target regions are highlighted in grey.

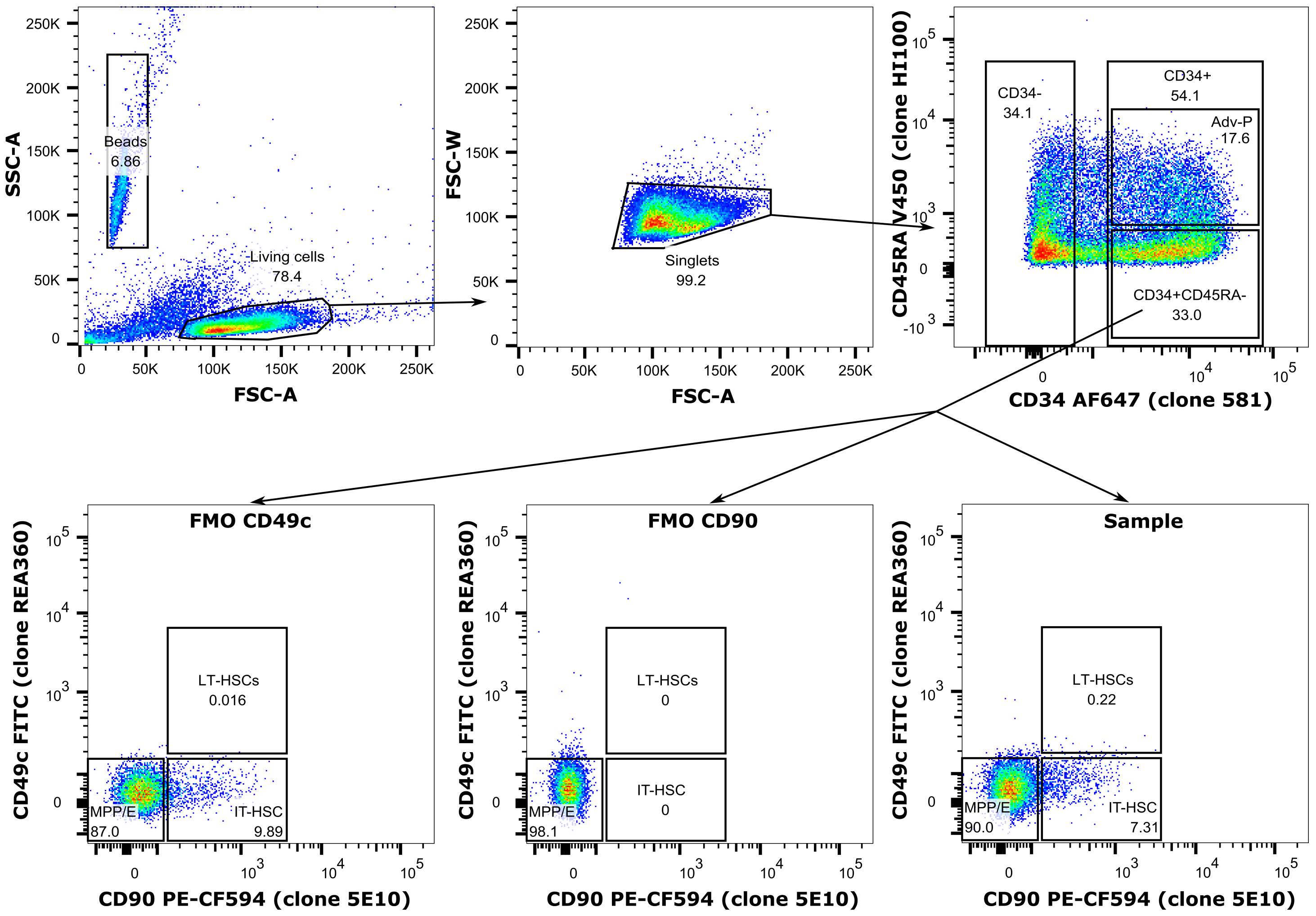

**Figure S6: Example gating hierarchy.** FMO stands for full-minus-one, where the indicated antibody was left out of the panel.
